# Post-ischemic inactivation of HIF prolyl hydroxylases in endothelium promotes maladaptive kidney repair by inducing glycolysis

**DOI:** 10.1101/2023.10.03.560700

**Authors:** Ratnakar Tiwari, Rajni Sharma, Ganeshkumar Rajendran, Gabriella S. Borkowski, Si Young An, Michael Schonfeld, James O’Sullivan, Matthew J. Schipma, Yalu Zhou, Guillaume Courbon, Valentin David, Susan E. Quaggin, Edward Thorp, Navdeep S. Chandel, Pinelopi P. Kapitsinou

## Abstract

Ischemic acute kidney injury (AKI) is common in hospitalized patients and increases the risk for chronic kidney disease (CKD). Impaired endothelial cell (EC) functions are thought to contribute in AKI to CKD transition, but the underlying mechanisms remain unclear. Here, we identify a critical role for endothelial oxygen sensing prolyl hydroxylase domain (PHD) enzymes 1-3 in regulating post-ischemic kidney repair. In renal endothelium, we observed compartment-specific differences in the expression of the three PHD isoforms in both mice and humans. We found that post-ischemic concurrent inactivation of endothelial PHD1, PHD2, and PHD3 but not PHD2 alone promoted maladaptive kidney repair characterized by exacerbated tissue injury, fibrosis, and inflammation. Single-cell RNA-seq analysis of the post-ischemic endothelial PHD1, PHD2 and PHD3 deficient (*PHD^TiEC^*) kidney revealed an endothelial glycolytic transcriptional signature, also observed in human kidneys with severe AKI. This metabolic program was coupled to upregulation of the *SLC16A3* gene encoding the lactate exporter monocarboxylate transporter 4 (MCT4). Strikingly, treatment with the MCT4 inhibitor syrosingopine restored adaptive kidney repair in *PHD^TiEC^* mice. Mechanistically, MCT4 inhibition suppressed pro-inflammatory EC activation reducing monocyte-endothelial cell interaction. Our findings suggest avenues for halting AKI to CKD transition based on selectively targeting the endothelial hypoxia-driven glycolysis/MCT4 axis.

## INTRODUCTION

Ischemia is the most common cause of acute kidney injury (AKI), a condition that affects approximately 10-15% of all hospitalized patients, and more than 50% of critically ill patients (1-3). Failure to fully recover from AKI may lead to persistent functional impairment and development of chronic and progressive kidney disease, leading to end stage renal disease (ESRD) (4). Given the lack of treatments that successfully promote kidney repair, it is crucial to better understand the mechanisms that regulate AKI to chronic kidney disease (CKD) transition and identify novel therapeutic targets.

Severe or prolonged ischemic injury can overwhelm the kidney’s reparative ability and trigger a cascade of detrimental responses within the renal microenvironment. Tubular epithelial cells undergo dedifferentiation, apoptosis, and impaired regeneration, leading to the disruption of the nephron structure (5). Immune cells, such as neutrophils and macrophages, infiltrate the injured tissue and perpetuate parenchymal cell injury and fibrosis (6-8). Fibroblasts become hyperactive, excessively depositing collagen and promoting tissue scarring (7, 9). In this milieu, endothelial cells (ECs) form a metabolically dynamic barrier that when impaired promotes immune cell migration, dysregulated vascular tone, and permeability (10, 11). As a result, the reduced blood flow limits oxygen delivery to affected areas leading to hypoxia. Besides the microvasculature dysfunction, the induction of hypoxia in the post-ischemic kidney involves multiple factors such as increased oxygen consumption, mitochondrial dysfunction, reduced oxygen carrying capacity due to anemia, and hindered oxygen diffusion by extracellular matrix (ECM) buildup (12). While hypoxia has been linked to CKD progression, the underlying cellular mechanisms remain poorly understood.

Being in direct contact with the blood, ECs are well equipped to detect changes in oxygen levels through prolyl-4-hydroxylase domain-containing proteins (PHDs, also known as EGLNs), which control the stability of HIF-α. In the presence of oxygen, PHDs hydroxylate HIF-α subunits at highly conserved proline residues promoting their ubiquitination by the VHL (von-Hippel–Lindau) enzyme and ultimate proteasomal degradation (13). Under hypoxia, the reduced catalytic activity of PHDs results in HIF-α stabilization. HIF-α then translocates to the nucleus, where it dimerizes with the constitutively expressed HIF-β subunit (also known as ARNT), regulating the transcription of a broad array of genes. Genes involved in survival, metabolism, and angiogenic activity of vascular ECs are regulated by HIF with major implications in controlling vascular development and disease settings such as cancer and inflammatory processes (14-17). While PHD2 is considered the main oxygen sensor regulating HIF-α protein levels in normoxia, other PHD isoforms contribute to HIF regulation in particular cell types or conditions, dependent on their abundance (6, 18). In vascular endothelium, significant heterogeneity across organs introduces an additional layer of complexity. For instance, we and others have shown a crucial role for PHD2 in regulating endothelial HIF-α in the lung, and endothelial PHD2 inactivation led to pulmonary hypertension (19, 20). On the other hand, our recent studies showed decreased responsiveness of the kidney endothelial HIF-α to PHD2 loss alone (21). Indeed, the expression of other PHD isoforms in the renal vascular bed as demonstrated by single-cell RNA sequencing (scRNA-seq) data implied their potential contribution in regulating endothelial HIF activation in the kidney (21). Due to this complexity of the PHD/HIF system in vascular endothelium, their relevance in kidney repair after ischemic injury remain elusive.

To delineate the role of endothelial oxygen sensing in regulating post-ischemic kidney repair, we have generated a set of conditional mouse strains in which we induced endothelial inactivation of PHD2 alone, or in combination with PHD1 and PHD3 taking advantage of the *Cdh5Cre(PAC)ER* transgenic mice (21, 22). While PHD2 inactivation alone did not alter kidney repair, simultaneous post-ischemic inactivation of endothelial PHD1, PHD2 and PHD3 induced robust HIF activation and exacerbated fibrosis, inflammation, and capillary dropout. ScRNA-seq analysis revealed a signature significant for endothelial hypoxia and glycolysis in association with proinflammatory responses. Finally, using a pharmacologic approach, we identified the hypoxia-regulated lactate exporter monocarboxylate transporter 4 (MCT4) as a potential target to suppress maladaptive pro-inflammatory responses and halt AKI to CKD transition.

## RESULTS

### Post-ischemic inactivation of endothelial PHD2 does not affect post-ischemic kidney injury

Our prior studies have shown that constitutive or acute inactivation of PHD2 in ECs prior to kidney ischemia-reperfusion injury (IRI) prevents post-ischemic injury and inflammation (21, 23). While these findings show the beneficial effects of endothelial PHD/HIF signaling to promote kidney resilience against ischemic insult, they cannot be extrapolated to kidney repair following established injury, in which distinct responses control regeneration of the damaged tissue (24, 25). To ask whether inhibition of endothelial PHD2 regulates kidney repair, we took advantage of tamoxifen-inducible *Cdh5-CreER^T2^;Phd2^f/f^*mice (*PHD2^iEC^*) (21, 22), which allow induction of recombination following established kidney injury. We first assessed the endothelial recombination efficiency achieved when tamoxifen treatment started on day 1 post renal unilateral IRI (uIRI) and included 4 total doses given every other day, as indicated in Supplemental Figure 1A. Using *Cdh5(PAC)CreER^T2^-Rosa26-mTmG* mice (21), which allow tracking of recombinant cells by GFP expression, we performed flow cytometry (Supplemental Figure 1B) and found that ∼100 % of GFP^+ve^ cells in day 14 post-ischemic kidneys were also CD31^+ve^. This recombination efficiency was comparable to the contralateral kidneys. Otherwise, at day 14 post-IRI, there was a ∼50% reduction in kidney ECs, which is expected in the setting of IRI-induced capillary rarefaction (Supplemental Figure 1C).

After validating the recombination efficiency of the *Cdh5(PAC)CreER^T2^*system in the post-ischemic kidney, adult male *PHD2^iEC^* mice and their *Cre^-^* littermates were subjected to uIRI, followed by four tamoxifen injections and analysis on day 14, as indicated in Figure 1A. Assessment of tubular injury on H&E-stained sections of day 14 post-ischemic kidneys revealed that *PHD2^iEC^*had comparable tubular damage to *Cre^-^* littermate controls. Furthermore, quantitative analysis of Picro-Sirius red staining showed no significant difference in collagen accumulation between mutants and controls (Figure 1B). Finally, day 14 post-ischemic kidneys showed significantly increased transcripts of profibrotic genes lysyl oxidase-like 2 (*Loxl2*), transforming growth factor beta 1 (*Tgfb1*), and smooth muscle cell a-actin (*Acta2*) compared to their corresponding contralateral, but there was no significant change between the two genotypes (Figure 1C). Therefore, acute post-ischemic endothelial inactivation of PHD2 does not affect kidney repair and transition to CKD following IRI.

**Figure 1.**
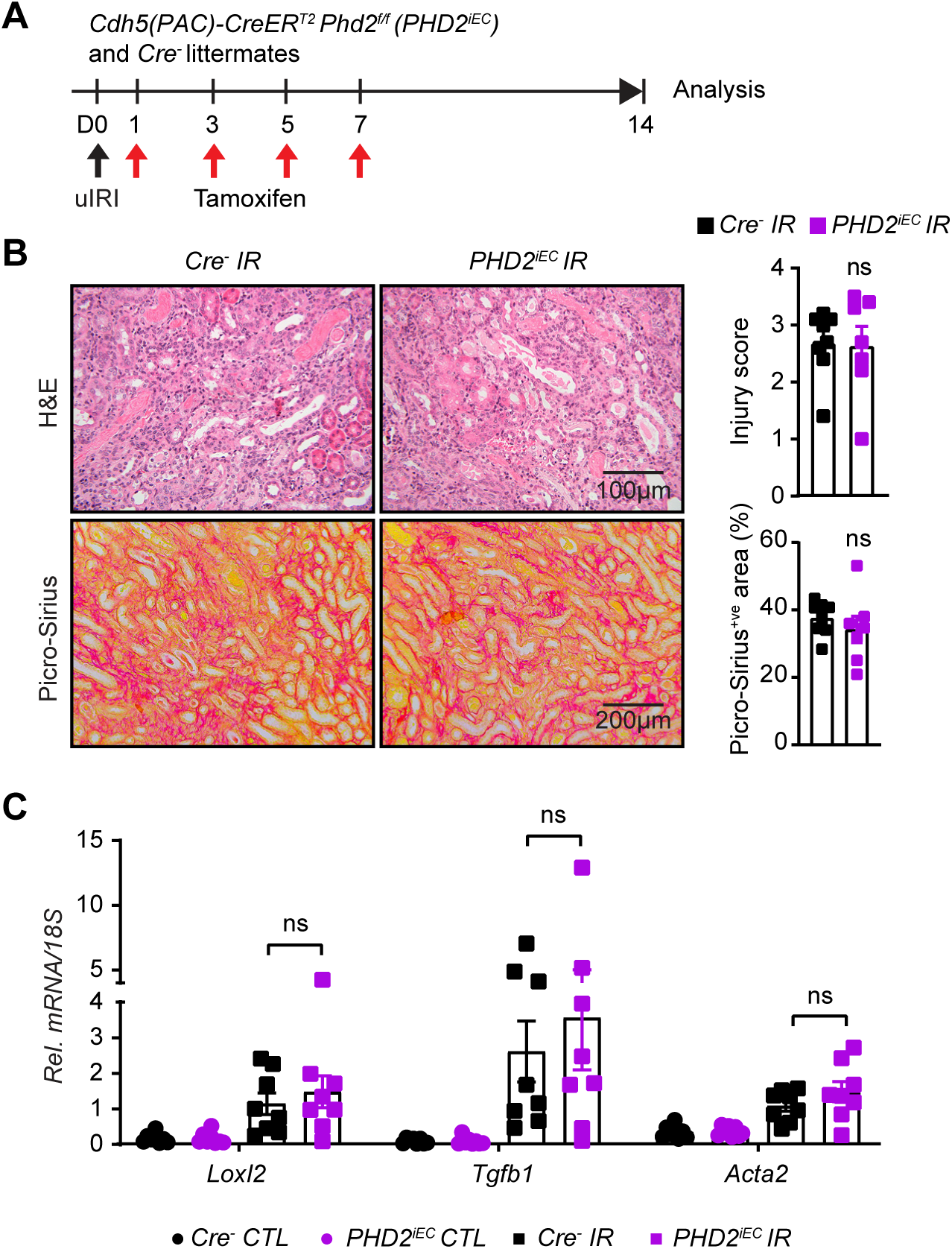
Post-ischemic inactivation of endothelial PHD2 does not alter post-ischemic kidney injury. (**A**) Experimental scheme illustrates the timing of unilateral renal artery clamping, tamoxifen administration and analysis. (**B**) Representative images of H&E and Picro-Sirius red stained sections from day 14 post-ischemic kidneys of *PHD2^iEC^* mutants and their *Cre^-^* littermates. Right panels show tubular injury score (Top) and semi-quantitative analysis of Picro-Sirius red^+ve^ area in the indicated geno-types. Scale bars indicate 100 μm and 200 μm for H&E and Picro-Sirius red images, respectively. (**C**) mRNA levels of *Loxl2*, *Tgfb1* and *Acta2* in IR and CTL kidneys from *PHD2^iEC^* mice and their *Cre^-^* controls at day 14 after unilateral IRI. All bars show mean ± SEM. For (**B**), unpaired t-test with Welch’s correction was used. For (**C**), statistics were determined using one-way ANOVA with Sidak correction for multiple comparisons. *, *P* <0.05; ns, not statistically significant. uIRI, unilateral ischemia reperfusion injury; CTL, contralateral; IR, kidney subjected to uIRI; Rel., relative.

### Kidney endothelium shows compartment-specific differential expression of PHD1, 2, and 3 in mice and humans

We recently reported that inactivation of endothelial PHD2 alone was not sufficient to stabilize HIF in the renal endothelium (21), raising the possibility for significant contributions of PHD1 and/or PHD3 in regulating renal endothelial HIF protein levels. Indeed, while PHD2 is considered the primary regulator of HIF-α stability under normal conditions (26-28), several studies have suggested a significant role for other PHD isoforms in controlling HIF-α activity under different circumstances depending on differential induction and cell type specific PHD expression (6, 18). Nevertheless, the function of PHD isoforms in different vascular beds remains unexplored. To better understand the role of oxygen sensors in renal vasculature, we took advantage of publicly available scRNA-seq data and characterized the expression of different *Phd* isoforms in kidney EC. We first analyzed the expression of *Phd1* (*Egln2), Phd2 (Egln1)*, and *Phd3 (Egln3)* in scRNA-seq data of renal ECs (RECs) extracted from the murine EC Atlas database (Figure 2A-C) (29). We found that *Phd1* was ubiquitously expressed, whereas *Phd2* and *Phd3* were mainly expressed in cortical and medullary RECs. Glomerular RECs showed particularly low expression for *Phd3*. To further validate the differential expression of PHDs in murine kidney endothelium, we immunostained kidney sections from tamoxifen treated *Cdh5(PAC)CreER^T2^-Rosa26-mTmG* reporter mice with antibodies against PHD1, PHD2, and PHD3. Immunolabelling confirmed ubiquitous expression of PHD1, expression of PHD2 in cortical and medullary RECs, and low expression of PHD3 in glomerular RECs (Supplemental Figure 2).

**Figure 2.**
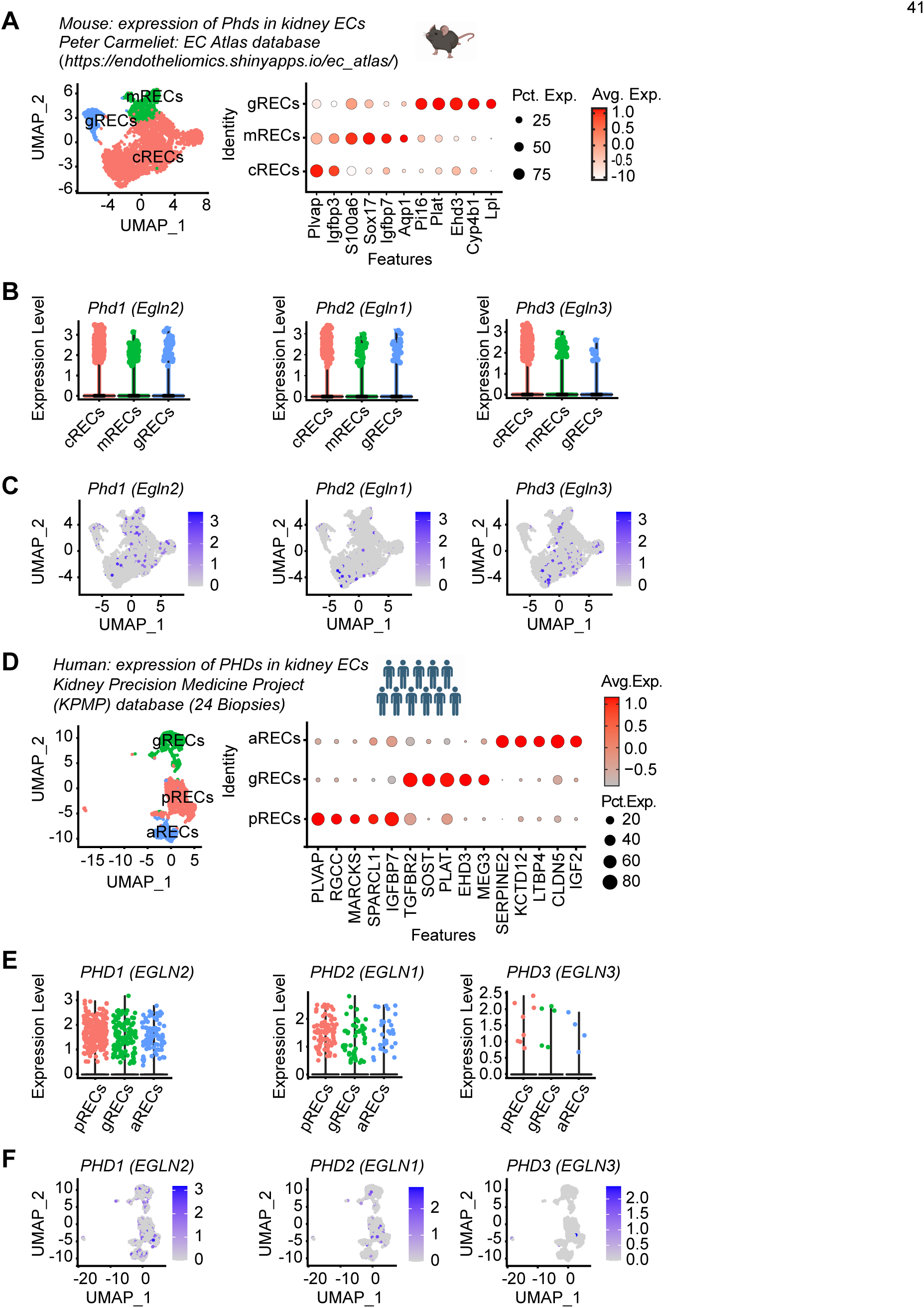
scRNA-seq analysis shows differential expression of *PHD1*, *PHD2* and *PHD3* in kidney ECs in mouse and humans. (**A-C**) scRNA-seq analysis of RECs extracted from mouse EC Atlas database (https://endotheliomics.shinyapps.io/ec_atlas/). (**A**) Uniform manifold approximation and projection (UMAP) plot shows three EC clusters; cortical RECs (cRECs), medullary RECs (mRECs) and glomerular RECs (gRECs). Dot plot displays gene expression patterns of cluster-enriched markers. Violin plots (**B**) and feature plots (**C**) show the expression of *Phd1* (*Egln2*), *Phd2* (*Egln1*) and *Phd3* (*Egln3*) in murine RECs. (**D-F**) scRNA-seq analysis of RECs extracted from normal human kidney biopsies (n=24). (**D**) UMAP plot shows arteriolar RECs (aRECs), glomerular RECs (gRECs) and peritubular RECs (pRECs). Dot plot illustrates gene expression patterns of cluster-enriched markers. Violin plots (**E**) and feature plots (**F**) show the expression of *PHDs* in different RECs clusters.

Next, we sought to characterize the expression of PHD isoforms within human kidney endothelium. To this end, we analyzed scRNA-seq data of human kidneys available through KPMP (Figure 2D-F and Supplemental Table 1) (30). ECs were identified by the expression of CD34 and endomucin (EMCN) and were clustered to three sub-clusters based on the expression of previously validated specific markers; *PLVAP* in peritubular ECs, *EHD3* in glomerular and *SERPINE2* in arteriolar ECs (Figure 2D) (30). This analysis demonstrated that *PHD1* was the most highly expressed isoform whereas *PHD3* showed the lowest expression. Among the different compartments, PHD2 was predominantly expressed in peritubular capillary ECs (Figure 2E and F). Together these studies suggest endothelial compartment-specific differences in the expression of the three PHD isoforms in mice and humans.

### Post-ischemic concurrent inactivation of endothelial PHD1, 2 and 3 promotes HIF activation and maladaptive kidney repair

Because we found that all PHD isoforms are expressed in kidney endothelium with variable abundance within different compartments, we next generated mice that allow acute endothelial specific inactivation of PHD1, PHD2 and PHD3 (*Cdh5(PAC)CreER^T2^; Phd1^f/f^ Phd2^f/f^ Phd3^f/f^ mice*; *PHD^TiEC^*) and predicted that the resulting mutants will show HIF stabilization in endothelium. One week after the completion of tamoxifen treatment (Supplemental Figure 3A), co-immunostaining with antibodies against PHD1, 2 and 3 and EMCN demonstrated successful ablation of PHD1, 2 and 3 in kidney ECs of *PHD^TiEC^*compared to *Cre^-^* littermates (Supplemental Figure 3B). Further, immunoblot assessment of kidney nuclear extracts isolated from *PHD^TiEC^*and *Cre^-^* littermates revealed significant stabilization of HIF-1α and HIF-2α in the kidneys of *PHD^TiEC^*mice (Supplemental Figure 3C). Baseline histopathological analysis showed normal kidney morphology without fibrosis in *PHD^TiEC^* mice as indicated by H&E and Picro-Sirius red staining (Supplemental Figure 3D).

To determine the impact of post-ischemic inactivation of endothelial PHD1, PHD2 and PHD3 on kidney repair following IRI, *Cre^-^* and *PHD^TiEC^* adult mice were subjected to unilateral renal artery clamping for 25 minutes followed by tamoxifen treatment and analysis on day 14 (Figure 3A). Notably, histological analysis of day 14 post-ischemic kidneys revealed higher tubular dilatation and cast formation in *PHD^TiEC^* mice than in *Cre^-^* controls (Figure 3B). Furthermore, day 14 post-ischemic kidneys from *PHD^TiEC^*mutants showed ∼38% increase in collagen deposition compared to *Cre^-^* littermates as indicated by Picro-Sirius red staining (n=6-8, *P* <0.05) (Figure 3B). Accordingly, qPCR analysis also showed significant transcriptional induction of pro-fibrotic genes *Loxl2* (P <0.05), *Tgfb1* (*P* <0.01), and *Acta2* (*P* <0.05) indicating increased post-ischemic kidney fibrosis in *PHD^TiEC^*mutants compared to *Cre^-^* controls (Figure 3C). Because peritubular capillary rarefaction is a hallmark of fibrotic kidneys, we also assessed capillary density by EMCN staining and found that EMCN^+ve^ area was reduced by 35% in day 14 post-ischemic kidneys from *PHD^TiEC^* mice (n= 4, *P*<0.05) (Figure 3D) (31-34).

**Figure 3.**
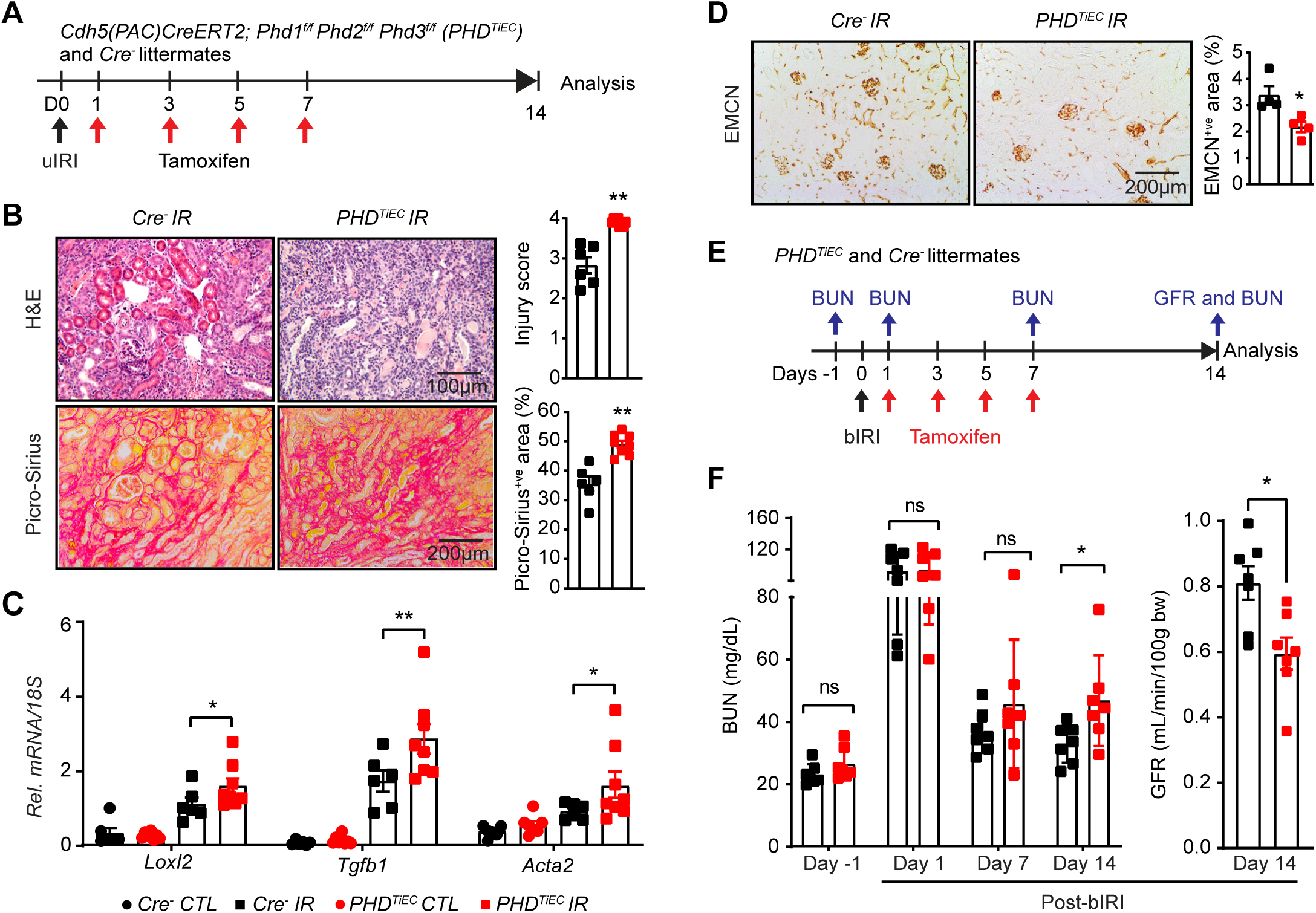
Post-ischemic simultaneous inactivation of endothelial PHD1, 2, and 3 promotes maladaptive kidney repair. (**A**) Scheme illustrating the experimental strategy applied for unilateral renal IRI (uIRI) studies. *PHD^TiEC^* mice and their *Cre^-^* littermates were subjected to 25 minutes of unilateral renal artery clamping. Treatment with tamoxifen was started on day 1 post uIRI involving 4 IP doses given every other day. Mice were sacrificed for histopathological and molecular analysis on day 14 post uIRI. (**B**) Representative images of H&E and Picro-Sirius red stained sections as well as tubular injury score and semi-quantitative analysis of Picro-Sirius red^+ve^ area on day 14 post-ischemic kidneys from *PHD^TiEC^* mice and *Cre^-^* littermates. Scale bars indicate 100 μm and 200 μm for H&E and Picro-Sirius red images, respectively. (**C**) mRNA levels of *Loxl2*, *Tgfb1* and *Acta2* in IR and CTL kidneys from *PHD^TiEC^* mice or their *Cre^-^* controls at day 14 after uIRI. (**D**) Representative images of EMCN immunostaining and semiquantitative analysis of EMCN^+ve^ peritubular capillary area on day 14 post-ischemic kidneys from *PHD^TiEC^* mice and *Cre^-^* littermates. (**E**) Scheme depicting the experimental workflow for bilateral renal IRI (bIRI) studies. *PHD^TiEC^* mice and their *Cre^-^* littermates were subjected to 23 minutes of bilateral renal artery clamping followed by tamoxifen treatment as described in **A**. Serum BUN levels were measured on 1 day prior to bIRI (baseline) and on days 1, 7 and 14 post bIRI. GFR was measured by FITC-sinistrin clearance on day 14 post bIRI using the MediBeacon transdermal GFR monitor system. (**F**) Serum BUN levels at different time points and GFR measurements on day 14 post bIRI. All bars show mean ± SEM of each group (n=4-8). For (**B**), (**D**) and (**F**), statistics were determined by unpaired t-test with Welch’s correction. For (**C**), one-way ANOVA with Sidak correction for multiple comparisons was used. *, *P*< 0.05; **, *P*< 0.01; ns, not statistically significant. CTL, contralateral kidney; IR, kidney subjected to IRI; bIRI, bilateral IRI; Rel., relative.

To assess the impact of post-ischemic endothelial inactivation of *Phd1, 2,* and *3* on recovery of kidney function after IRI, *PHD^TiEC^* and *Cre^-^* littermate mice were subjected to bilateral renal IRI (bIRI), followed by tamoxifen treatment (Figure 3E). Blood urea nitrogen (BUN) levels were measured one day prior to surgery (baseline) and on days 1, 7 and 14 after bIRI surgery. As expected, on day 1 after bIRI, BUN levels increased to similar degree for both genotypes without a significant difference on day 7. However, on day 14 post bIRI, *PHD^TiEC^* had significantly increased BUN levels by 43% compared to *Cre^-^* mice (*P*<0.05). In accordance, GFR measurements on day 14 post bIRI showed 26% reduction in GFR of *PHD^TiEC^* mice compared to their *Cre^-^* littermates (n =7, *P*<0.01) (Figure 3F). In conclusion, our findings demonstrate that EC-specific inactivation of PHD1, PHD2 and PHD3 after IRI promotes maladaptive kidney repair characterized by tubular injury, interstitial fibrosis and capillary drop-out; these changes lead to impaired recovery of kidney function.

### scRNA-seq reveals distinct cell-specific responses in post-ischemic PHD^TiEC^ mutant kidney

To investigate the impact of post-ischemic endothelial PHD inactivation on cell-type specific alterations in the context of kidney repair following IRI, we carried out scRNA-seq on day 14 post-ischemic kidneys from *PHD^TiEC^*and *Cre^-^* control mice. Using the 10X Genomics Chromium platform, we aimed for 10,000 cells per group (Figure 4A). After quality filtering, a total of 16,698 (7,955 from *Cre^-^* and 8,743 from *PHD^TiEC^*) cells remained for subsequent analysis. We identified 28 different cell clusters using unsupervised graph-based clustering followed by principal component analysis (PCA) and uniform manifold approximation and projection (UMAP) (Figure 4B). When we overlayed *Cre^-^* IR and *PHD^TiEC^* IR samples, we observed similar cell clusters in each genotype (Supplemental Figure 4). Based on expression of reported canonical marker genes (Supplemental Table 2), we identified all expected kidney cell types; proximal tubule (PT), Injured PT, thick ascending limb (TAL), distal convoluted tubule (DCT), collecting duct (CD), proliferating CD (pCD), inner medullary collecting duct (IM-CD), collecting duct-intercalated cells (CD-IC), intercalated cells (IC), parietal cells (PAR), fibroblasts (FIB), pericytes (PER), endothelial cells 1-3 (EC1-3), urothelial cells (URO), Macrophages 1-4 (Mφ1-4), proliferating macrophages (pMφ), C1q (C1qa, C1qb and C1qc) expressing immune cells (C1q-IM), T cells (T), proliferating T cells (pT), natural killer cells (NK), B cells (B), dendritic cells (DEN), and neutrophils (NEU) (Figure 4C). Consistent with the aforementioned increased fibrosis, analysis of the relative cell proportions for each genotype showed increased fibroblasts by ∼1.9-fold in the post-ischemic *PHD^TiEC^* kidney compared to control. Next, we performed differential gene expression analysis for tubular clusters between the two genotypes. After applying cutoff criteria of a log_2_ fold change >0.2 and adjusted *P*<0.05, PT, Inj-PT, TAL, DCT, CD, and CD-IC clusters of *PHD^TiEC^* IR kidney showed 163, 90, 209, 269, 211 and 250 differentially expressed genes (DEGs), respectively (Supplemental Datasheet 1). Gene set enrichment analysis (GSEA) demonstrated significant differences in tubular cells between *PHD^TiEC^* and *Cre^-^*. Within the tubular clusters, the Top Hallmark gene signatures were TNFA signalling via NFkB and Hypoxia pathways for PT; Apoptosis, TNFA signalling via NFkB, and Hypoxia pathway for DCT; TNFA signalling via NFkB, Apoptosis, and Hypoxia pathway for CD (Figure 4D & Supplemental Datasheet 2). Overall, these results show that the exacerbated fibrotic response in post-ischemic kidneys of *PHD^TiEC^* mutants is associated with a pro-inflammatory signature in tubular cells.

**Figure 4.**
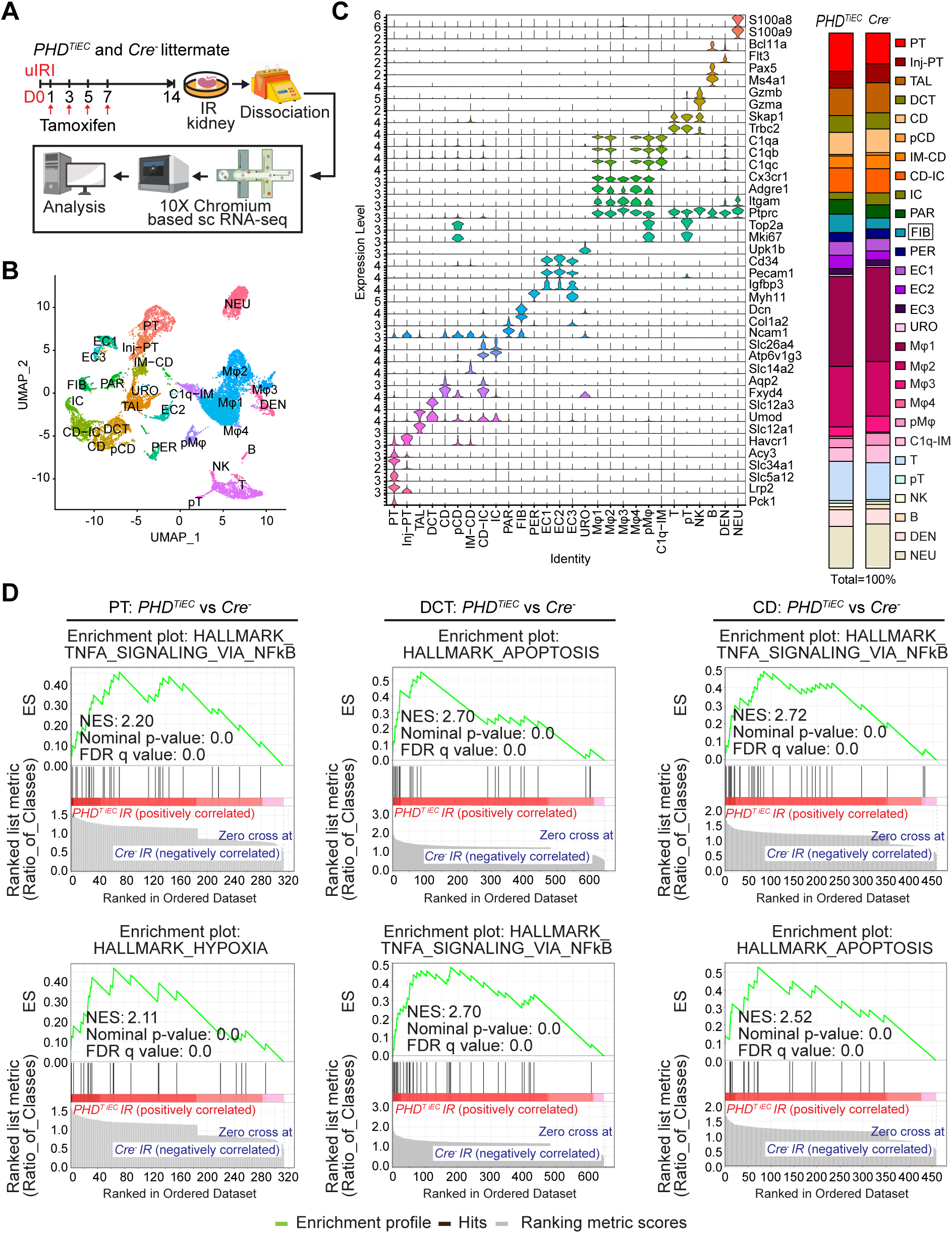
scRNA-seq analysis reveals the cellular landscape of day 14 post-ischemic kidneys of *PHD^TiEC^* and control mice. (**A**) Scheme illustrating the experimental strategy applied for scRNA-seq analysis. A *PHD^TiEC^* mouse and a *Cre^-^* littermate were subjected to 25 minutes of unilateral renal artery clamping. Treatment with tamoxifen was started on day 1 post uIRI and involved 4 IP doses given every other day. Mice were sacrificed for scRNA-seq analysis on day 14 post uIRI. IR kidneys were isolated and single cell suspension was prepared and used for scRNA-seq analysis (n=1 per genotype). (**B**) Uniform manifold approximation and projection (UMAP) plot representation of the cell classification in day 14 post-ischemic kidneys from *PHD^TiEC^* and *Cre^-^* mice. (**C**) Violin plots display characteristic marker genes for each identified cell population. Right side bar plot shows cell proportions in day 14 post-ischemic kidneys from *Cre^-^* and *PHD^TiEC^* mice. Highlighted is the increased proportion of fibroblasts in *PHD^TiEC^* post-ischemic kidney compared to *Cre^-^* control. Proximal tubule (PT), Injured proximal tubule (Injured-PT), Thick ascending limb (TAL), Distal convoluted tubule (DCT), Collecting duct (CD), Proliferating CD (pCD), Inner medullary collecting duct (IM-CD), Collecting duct-intercalated cells (CD-IC), Intercalated cells (IC), Parietal cells (PAR), Fibroblasts (FIB), Pericytes (PER), Endothelial cells 1-3 (EC1-3), Urothelial cells (URO), Macrophages 1-4 (Mφ1-4), Proliferating macrophages (pMφ), C1q (C1qa, C1qb and C1qc) expressing immune cells (C1q-IM), T cells (T), Proliferating T cells (pT), Natural killer cells (NK), B cells (B), Dendritic cells (DEN), and Neutrophils (NEU). (**D**) Top 2 enriched Hallmark pathways emerged in GSEA (gene set enrichment analysis) hallmark analysis of differentially expressed genes (DEGs) for PT, DCT and CD clusters of *PHD^TiEC^* post-ischemic kidney as compared to *Cre^-^* control.

### Medullary renal ECs of post-ischemic PHD^TiEC^ mutant kidney show increased glycolysis

Next, we focused on the three EC clusters annotated as medullary RECs (mRECs), cortical Renal ECs (cRECs), and RECs expressing mesenchymal markers (EndMT-RECs) based on marker genes established from the Carmeliet group (35) and others (36-38) (Figure 5A, Supplemental Figure 5 and Supplemental Figure 6). Among the EC clusters, mRECs showed the majority of DEGs between the two genotypes (58 genes with adjusted *P* value <0.05) (Supplemental datasheet 3), while the numbers of DEGs for cRECs and EndMT-RECs were more limited (2 and 12 genes respectively). GSEA analysis of DEGs in mRECs identified Hypoxia and Glycolysis as the top two significantly enriched Hallmark pathways followed by MTORC1 signaling and Epithelial Mesenchymal Transition pathways in *PHD^TiEC^* compared to *Cre^-^* mouse (Figure 5B and Supplemental datasheet 3). Consistent with enhanced glycolytic activity, the glycolysis related genes solute carrier family 2 (facilitated glucose transporter), member 1 (*Scl2a1*), glucose-6-phosphate isomerase 1 (*Gpi1*), phosphofructokinase, liver, B-type (*Pfkl*), aldolase A, fructose-bisphosphate (*Aldoa*), triosephosphate isomerase 1 (*Tpi1*), phosphoglycerate kinase 1 (*Pgk1*), enolase 1, alpha non-neuron (*Eno1*), pyruvate kinase, muscle (*Pkm*), lactate dehydrogenase A (*Ldha*) and solute carrier family 16, member 3 (*Slc16a3*) were significantly upregulated in mRECs of *PHD^TiEC^* IR kidney compared to *Cre^-^* control (Figure 5C).

**Figure 5.**
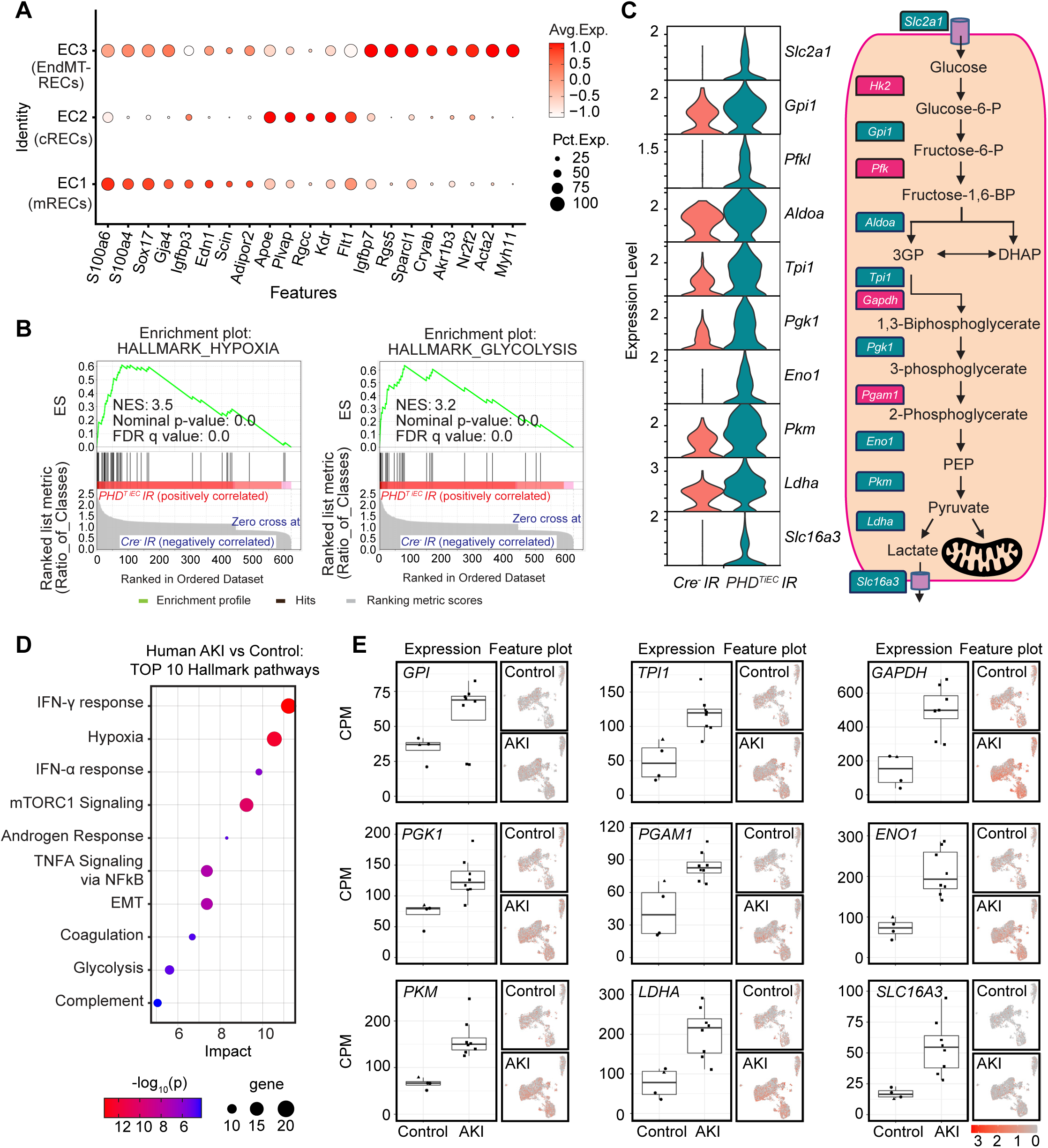
Post-ischemic endothelial PHD inactivation induces a hypoxia and glycolysis gene signature in mRECs. (**A**) Dot plot visualization shows the expression of marker genes used to identify cRECs, mRECs and EndMT-RECs clusters. (**B**) GSEA in mRECs of *PHD^TiEC^* kidney compared to control. Among the most highly enriched Hallmark pathways were Hypoxia and Glycolysis. (**C**) Violin plots show significantly upregulated glycolytic genes in mRECs of *PHD^TiEC^* compared to control. Pathway diagram summarizes the functions of up-regulated genes (marked by teal boxes) in glycolysis. (**D-E**) snRNA-seq analysis of human kidney tissue from patients with severe AKI and controls (n=6-8). Analysis was performed on publicly available snRNA-seq data from Christian Hinze et al (39). (**D**) Bubble chart for top 10 enriched Hallmark pathways of upregulated DEGs in kidney ECs from patients with severe AKI compared to controls. (**E**) Box plots show the expression of glycolytic genes in kidney ECs in controls vs AKI patients. The expression levels of glycolytic genes were extracted from the online interface provided by Christian Hinze et al (https://shiny.mdc-berlin.de/humAKI). CPM, normalized counts per million; cRECs, cortical renal ECs; mRECs, medullary renal ECs; EndMT-RECs, renal ECs expressing mesenchymal markers.

To examine whether this metabolic profile in ECs was relevant to human AKI, we analysed publicly available single nucleus RNA sequencing (snRNA-seq) data from kidney tissues of patients with severe AKI (39). Hallmark analysis showed Hypoxia and Glycolysis among the top enriched pathways in ECs of human kidneys with AKI (Figure 5D and Supplemental datasheet 4). Specifically, we found upregulation of glucose-6-phosphate isomerase (*GPI*), triosephosphate isomerase 1 (*TPI1*), glyceraldehyde-3-phosphate dehydrogenase (*GAPDH*), phosphoglycerate kinase 1 (*PGK1*), phosphoglycerate mutase 1 (*PGAM1*), enolase 1 (*ENO1*), pyruvate kinase M1/2 (*PKM*), lactate dehydrogenase A (*LDHA*), and solute carrier family 16 member 3 (*SLC16A3*) in ECs from kidney tissues of patients with AKI compared to the control group (Figure 5E). Overall, these results demonstrate that post-ischemic endothelial inactivation of PHDs activates the hypoxia response leading to increased endothelial glycolysis, a metabolic response that is also observed in patients with severe AKI.

### Post-ischemic endothelial inactivation of PHD1, PHD2 and PHD3 induces EC derived pro-inflammatory responses and exacerbates macrophage accumulation

GSEA analysis of DEGs is mRECs showed significant enrichment of Gene Ontology Biological Process (GOBP) gene sets of Leukocyte Migration and Myeloid Leukocyte Migration. These gene signatures were driven by upregulation of macrophage migration inhibitory factor (*Mif*), C-X-C motif chemokine ligand 12 (*Cxcl12*), FMS-like tyrosine kinase 1 (*Flt1*), basigin (*Bsg*), vascular endothelial growth factor A (*Vegfa*), serine (or cysteine) peptidase inhibitor, clade E, member 1 (*Serpine1*), CD74 antigen (*Cd74*), sphingosine-1-phosphate receptor 1 (*S1pr1),* intercellular adhesion molecule 1 (*Icam1*), and integrin beta 1 (*Itgb1*) in *PHD^TiEC^* IR kidney compared to control (Figure 6A and B). To characterize immune cell alterations in the post-ischemic kidney microenvironment in the context of endothelial PHD inactivation we performed flow cytometric analysis (Supplemental Figure 7). Because at day 14 post uIRI, there was an established maladaptive repair phenotype, we focused our analysis on day 8, an earlier time point that may reveal changes in immune responses contributing to the observed phenotype. Notably, day 8 post-ischemic kidneys of *PHD^TiEC^*showed a 2.9-fold increase in macrophages (CD45^+^ CD11b^+^ F4/80^+^) (Figure 6C and D) compared to controls. F4/80 immunostaining showed diffuse and variable distribution within the post-ischemic kidneys of *PHD^TiEC^* mutants (Figure 6D). Taken together, these data support that endothelial post-ischemic PHD inactivation promotes maladaptive inflammatory responses and increased macrophage infiltration.

**Figure 6.**
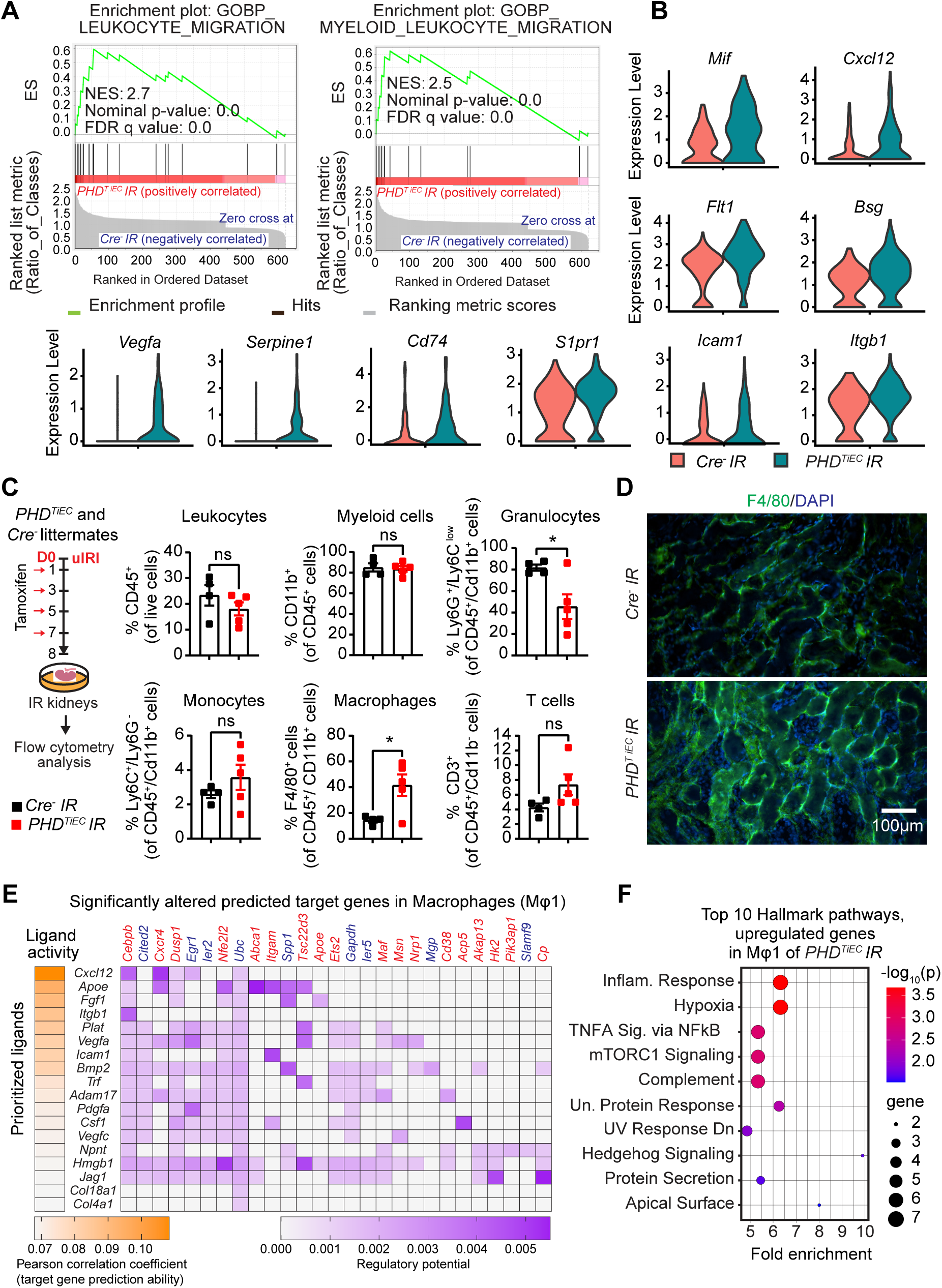
Post-ischemic inactivation of endothelial PHDs induces EC derived pro-inflammatory responses. (**A**) GSEA plots show significant enrichment for GO-biological processes (GOBP) of leukocyte migration and myeloid leukocyte migration in mRECs of *PHD^TiEC^* post-ischemic kidney compared to control. (**B**) Violin plots display the expression levels of pro-inflammatory genes associated with core enrichment in GOBP-leukocyte migration and myeloid leukocyte migration in mRECs of *PHD^TiEC^* compared to control. (**C**) Shown is the experimental strategy for flow cytometry analysis. Eight days after uIRI, post-ischemic kidneys from *PHD^TiEC^* and *Cre^-^* control mice were harvested, and flow cytometry analysis of immune cells was performed (n=4-5). Data are represented as mean ± SEM. Statistics were determined by unpaired t-test with Welch’s correction. (**D**) Representative images of immunofluorescence staining for F4/80 (green) and nuclear DAPI staining (blue) of day 8 post-ischemic kidneys from *PHD^TiEC^* and *Cre^-^* control mice. Images were captured using a Nikon Ti2 Widefield fluorescence microscope. Scale bar indicates 100 μm. (**E**) Shown is NicheNet analysis of mRECs communication with macrophage Mφ1 cluster in day 14 post-IRI kidney of *PHD^TiEC^* mutant compared to control. Top prioritized ligands expressed by mRECs (senders) and target genes that are significantly altered (red,upregulated genes; blue,downregulated genes) in the macrophages Mφ1 (receivers). The interaction pairs were derived from the NicheNet data sources and analysis. (**F**) Hallmark analysis of significantly upregulated genes in Mφ 1 cluster of post-ischemic *PHD^TiEC^* kidney compared to *Cre^-^* control. Top 10 pathways are shown. *, *P* <0.05; ns, not statistically significant.

To investigate how mRECs interact with macrophages, we employed NicheNet, a previously published algorithm, which utilizes expression data to infer the potential impact of ligand-receptor interactions on specific targets by integrating existing knowledge of signaling and regulatory networks (40). We assigned mRECs as “senders” and the different macrophage clusters as “receivers”. NicheNet analysis showed that ligands expressed by mRECs could explain 18 out of the 167 DEGs found in Mφ1 cluster of *PHD^TiEC^* mutant compared to control, while the impact was more limited for the other macrophage clusters (2/64 for Mφ2; 5/32 for Mφ3; Mφ4 and pMφ showed no interactions) (Figure 6E). Several notable ligands and target genes emerged from analysing the interaction of mRECs with Mφ1 cluster. One prominent example is the CXC chemokine CXCL12, which exhibited high expression in mRECs of *PHD^TiEC^* mutant. CXCL12 is a known HIF-target gene that can interact with the G proteincoupled chemokine receptor CXCR4, which is abundantly expressed in macrophages. This interaction has the potential to modulate macrophage activation contributing to pro-fibrotic function (41). Pathway analysis by GSEA identified several Hallmark pathways significantly enriched in Mφ1 cluster of *PHD^TiEC^* mutant. These pathways included Hypoxia, Inflammatory response, TNFA signaling via NFkB, mTORC1 signaling, Complement, and Unfolded Protein Response (Figure 6F). Taken together, these results suggest that post-ischemic endothelial PHD inactivation triggers an endothelial driven pro-inflammatory milieu with macrophages, which may contribute to tissue injury and fibrosis following kidney IRI.

### Treatment with the MCT4 inhibitor syrosingopine restores adaptive repair in post-ischemic kidney injury of PHD^TiEC^ mice

As the primary metabolic response of the post-ischemic endothelium to HIF stabilization in the context of PHD inactivation was increased glycolysis, we next focused on this aspect. While increased lactate generation and efflux via the transporter MCT4 represent the final steps of hypoxia-induced glycolysis (42-44), lactate export was recently shown to be among the four key steps controlling glycolytic flux (45). *Slc16a3*, the gene encoding MCT4 was significantly increased in mRECs of post-ischemic *PHD^TiEC^* kidney as well as in ECs of human kidneys with severe AKI. Indeed, immunofluorescence staining confirmed high expression of MCT4 protein in the post-ischemic kidney endothelium of *PHD^TiEC^* (Figure 7A and Supplemental Figure 8). We next sought to investigate whether increased endothelial MCT4 causally contributes to the maladaptive post-ischemic kidney repair observed in the *PHD^TiEC^* mice. To this end, we used syrosingopine, a dual inhibitor of monocarboxylate transporters MCT1 and MCT4 (60-fold higher potency on MCT4), which leads to end-product inhibition of glycolysis (46). As depicted in Figure 7B, *PHD^TiEC^* mice were treated with syrosingopine or vehicle starting on day 2 after uIRI and then every other day until day 14 post uIRI. Strikingly, we found that treatment with syrosingopine significantly suppressed tubular injury and fibrosis as indicated by ∼37% and ∼51 % reduction in injury scoring and collagen accumulation, respectively compared to the vehicle-treated mice (n=4-6, *P*<0.05) (Figure 7C and D). Furthermore, the mRNA levels of profibrotic genes *Loxl2*, *Tgfb1* and *Acta2* were significantly downregulated in syrosingopine treated group compared to vehicle treated group (n=5, *P<*0.05) (Figure 7E). Taken together, our studies demonstrate that inhibition of endothelial MCT4 by syrosingopine restores adaptive kidney repair following IRI in *PHD^TiEC^* mutants.

**Figure 7.**
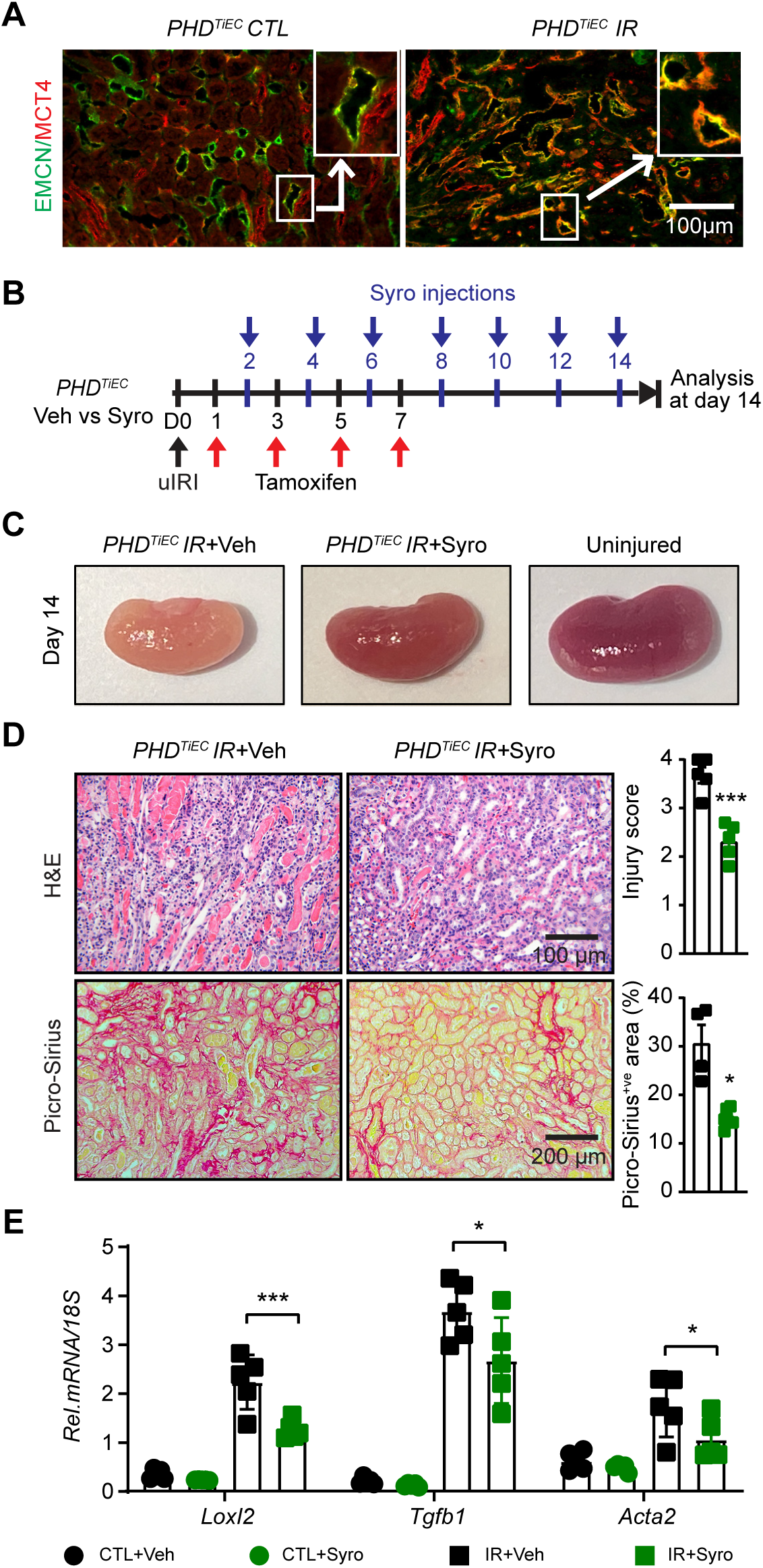
Post-IRI treatment with the MCT4 inhibitor syrosingopine restores adaptive kidney repair in *PHD^TiEC^* mice. (**A**) Repre-sentative images of immunofluorescence staining for MCT4 (red) and EMCN (green) of contralateral and day 14 post-ischemic kidneys from *PHD^TiEC^* mice. Zoom-in panels show the increased expression of MCT4 in EMCN^+ve^ cells in *PHD^TiEC^* post-ischemic kidney. Images were captured using a Nikon Ti2 Widefield fluorescence microscope. Scale bar, 100 μm. (**B**) Scheme shows the experimental protocol used. *PHD^TiEC^* mice were subjected to 25 minutes of unilateral renal artery clamping. Tamoxifen was started on day 1 post uIRI and was given every other day until day 7 post uIRI. Treatment with syrosingopine was started at day 2 post uIRI and was given every other day until day 14, when mice were sacrificed for histopathological and molecular analysis. (**C**) Representative images of uninjured kidney compared to day 14 post-ischemic kidneys treated with vehicle, or syrosingopine. All mice are *PHD^TiEC^* mutants. (**D**) Representative images of H&E and Picro-Sirius red stained day 14 post-ischemic kidneys from vehicle- vs syrosingopine-treated *PHD^TiEC^* mutants. Right: Tubular injury score and semiquantitative analysis of Picro-Sirius red^+ve^ area of day 14 post-ischemic kidneys for the indicated experimental groups. Scale bars indicate 100 μm and 200 μm for H&E and Picro-Sirius red images, respectively. (**E**) mRNA levels of *Loxl2*, *Tgfb1* and *Acta2* in CTL and IR kidneys from vehicle or syrosingopine-treated *PHD^TiEC^* mice on day 14 after uIRI. Data are represented as mean ± SEM. For (**D**), statistics were determined by unpaired t-test with Welch’s correction. For (**E**), statistics were determined using one-way ANOVA with Sidak correction for multiple comparisons. n=4-5; *, *P*< 0.05; ***, *P*< 0.001. uIRI, unilateral IRI; CTL, contralateral kidney; IR, kidney subjected to IRI; Veh, vehicle; Syro, syrosingopine.

### MCT4 inhibition diminishes endothelial derived pro-inflammatory responses induced by hypoxia-reoxygenation and IL-1β

To assess whether MCT4 inhibition acts cell-autonomously in ECs to suppress pro-inflammatory responses, we performed in vitro experiments in which human primary pulmonary artery endothelial cells (HPAECs) were activated by 18 hours exposure to hypoxia (0.5% O_2_), followed by reoxygenation for 8 hours in the presence of IL-1β treatment (1 ng/mL) (Figure 8A). To inhibit MCT4, we first used syrosingopine at concentration of 5 μΜ, which diminished lactate secretion, as indicated by reduction in extracellular lactate levels (Supplemental Figure 9A). We found that stimulation of HPAECs by hypoxia-reoxygenation and IL-1β led to ∼23- and ∼30-fold increase of vascular cell adhesion molecule 1 (*VCAM1*) and intercellular adhesion molecule 1 (*ICAM1*), respectively, while treatment with syrosingopine significantly suppressed the effects of hypoxia-reoxygenation and IL-1β on the expression of EC adhesion molecules (Figure 8B). Furthermore, under the same experimental conditions, treatment with syrosingopine significantly reduced adhesion of monocytes (THP-1 cells) to HPAECs by ∼48% (Figure 8C). Consistent responses were observed when we used siRNA against MCT4. With ∼ 67% knock down efficiency (Supplemental Figure 9B), we observed that *MCT4siRNA* significantly suppressed *VCAM1* and *ICAM1* transcripts in ECs stimulated by hypoxia-reoxygenation and IL-1β compared to negative control siRNA, reducing adhesion of monocytes to HPAECs by ∼40%. (Figure 8D and E). Taken together, these results demonstrate that MCT4 inhibition suppresses EC interaction with monocytes in a cell autonomous manner, an effect that may contribute to favorable effects exerted by syrosingopine, when given post kidney IRI.

**Figure 8.**
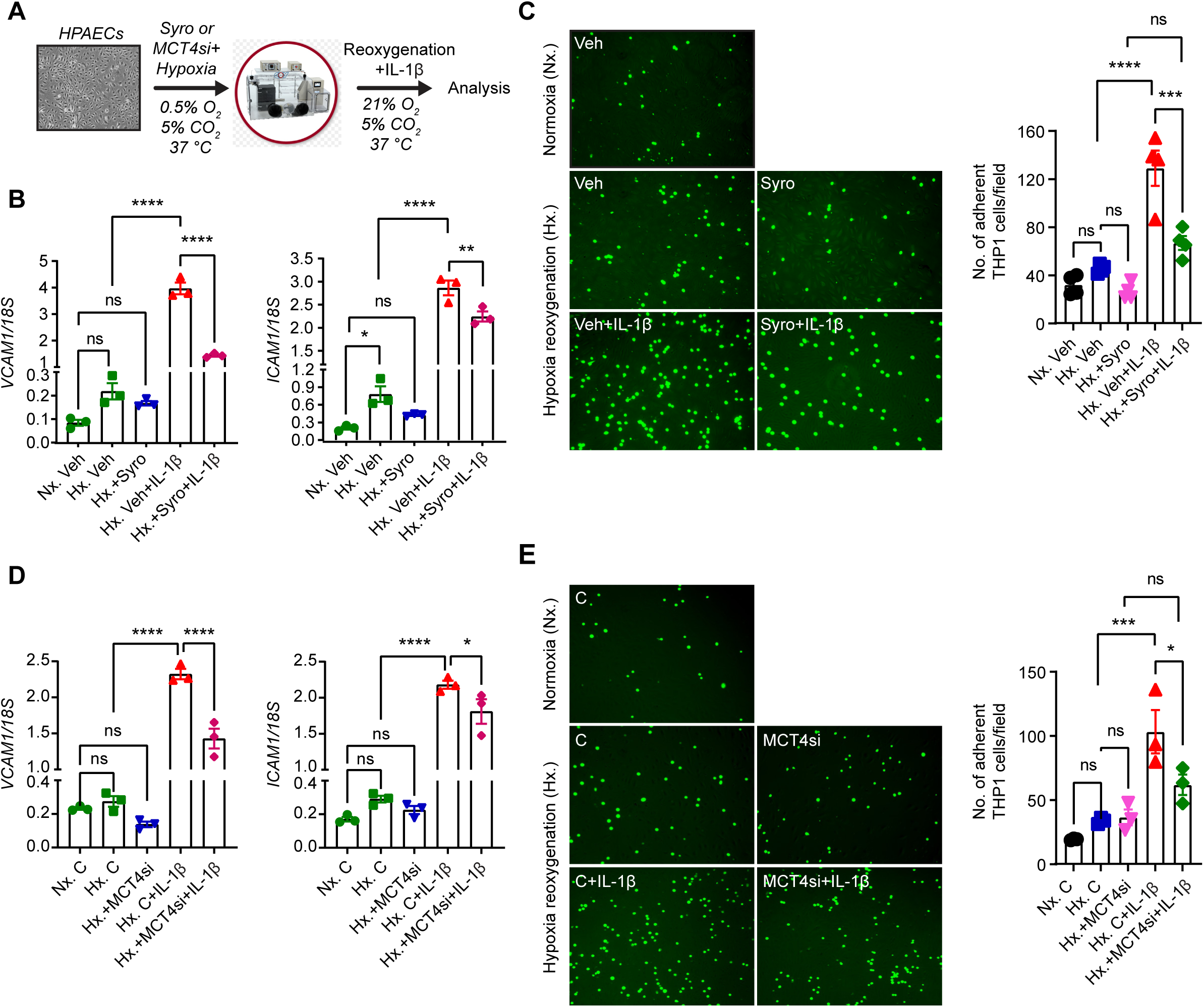
Syrosingopine or MCT4 knockdown suppresses the expression of EC adhesion molecules in HPAECs activated by Hypoxia/Re-oxygenation and IL-1β. (**A**) Experimental scheme for HPAECs subjected to 0.5% O_2_ for 18 hours in the presence of syrosingopine (5 µM) or MCT4 siRNA followed by reoxygenation for 8 hours in the presence of IL-1β (1 ng/ml). (**B**) mRNA levels of *VCAM1* and *ICAM1* in syrosingopine- vs vehicle-treated HPAECs, that were activated by Hypoxia/Reoxygenation and IL-1β. (**C**) THP1 monocyte adhesion to inflamed ECs. THP1 monocyte cells, labeled with Green CMFDA Dye, were introduced on a monolayer of HPAECs that had been subjected to the indicated experimental conditions. Following a 90-minute incubation period, floating cells were washed away and adhered THP1 cells were visualized using a fluorescent microscope and subsequently quantified. Representative images of fluorescent THP1 cells attached to ECs in different experimental groups are presented. Scale bar, 200 μm. (**D**) mRNA expression of *VCAM1* and *ICAM1* in HPAECs transfected with control or MCT4 siRNA and subjected to Hypoxia/Reoxygenation and IL-1β (**E**) THP1 monocyte adhesion to inflamed ECs. Labeled THP1 cells were introduced on a monolayer of HPAECs that had been subjected to the indicated experimental conditions. Following incubation, adhered THP1 cells were visualized quantified as in **C**. Representative images of fluorescent THP1 cells attached to ECs in different experimental conditions are presented. Scale bar, 200 μm. Data are represented as mean ± SEM. Statistics were determined by one-way ANOVA with Sidak correction for multiple comparisons. n=3-4; *, *P* < 0.05; **, *P* <0.01; ***, *P* <0.001; ****, *P* < 0.0001; ns, not statistically significant. Nx, Normoxia; Hx, Hypoxia/Reoxygenation; Veh, vehicle; Syro, syrosingopine; C, negative control siRNA; MCT4si, MCT4siRNA.

## DISCUSSION

In this study, we show that the regulation of endothelial oxygen sensing signaling has critical implications in post-ischemic kidney repair. Within the kidney endothelium, we demonstrate the expression of PHD1 and PHD3 isoforms along with PHD2 and show their critical contribution in regulating HIF signaling in the renal vascular bed. By investigating the outcomes of kidney repair in different conditional knock out strains, we provide evidence that concurrent loss of the three PHD isoforms and not PHD2 alone dictates post-ischemic kidney repair. We furthermore identify the endothelial glycolysis/MCT4 axis as a promising therapeutic target. Pharmacological inhibition of MCT4 holds the potential to interrupt the transition from AKI to CKD by mitigating inflammation.

PHD enzymes are non-heme iron-dependent dioxygenases that catalyze the four-electron reduction of O_2_. Specifically, they incorporate each O_2_ atom, respectively, into the tricarboxylic acid (TCA) cycle metabolite 2-oxoglutarate (2OG) and the substrate polypeptide, forming succinate, carbon dioxide, and *trans*-4-hydroxylated prolyl products (13). With *K*m values falling within the 230-250 μΜ range, PHDs are sensitive to fluctuations in local O_2_ in tissues, thus functioning as oxygen sensors (47). While PHD2 is considered the main oxygen sensor (13, 26), the other isoforms have been implicated in regulating hypoxia signaling (6). Consistently, we have demonstrated an overlapping pattern of expression for PHDs in kidney endothelium, which probably allows significant compensatory responses. This may explain the lack of HIF stabilization when endothelial PHD2 was individually deleted (21) in contrast to the robust HIF activation we observed when the three PHD isoforms were concurrently inactivated. While the regulation of HIF by PHDs in different vascular beds requires further investigation, our findings are particularly relevant for IRI, where significant hypoxia could limit the activity of all PHD isoforms. Furthermore, because PHDs are regulated not only by oxygen but also by factors such as 2OG, oxidative stress and abnormal concentrations of endogenous metabolites, PHD mediated hydroxylation can be altered even under normoxia, creating a “pseudo-hypoxic state” (13). For example, succinate, a TCA cycle metabolite known to accumulate with IRI can inhibit all three PHDs (48, 49). Therefore, various pathophysiological signals within the post-ischemic kidney microenvironment can inhibit endothelial PHDs dictating tissue remodeling. However, it remains unclear how PHDs in different cell types respond to hypoxic and pseudo-hypoxic signals and how the integrated responses alter kidney repair. Although it may be challenging, additional genetic studies involving inducible deletion of PHDs in different cell types are required to define the role of PHD/HIF pathway in the context of kidney repair.

Capillary rarefaction was exacerbated after ischemic kidney injury in the setting of postischemic endothelial PHD inactivation and was associated with the presence of partially “dedifferentiated” ECs as reflected by the EC-EndMT cluster revealed by ScRNA-seq. EndMT could be induced directly by HIF activation in the context of endothelial PHD inactivation as indicated by previous studies linking HIFs to EndMT in pulmonary and cardiac fibrosis by upregulating the transcription factor SNAI1 (50, 51). While the contribution of EndMT in our model is unclear, impaired angiogenic capacity could represent another EC autonomous mechanism by which PHD inactivation leads to capillary drop-out, as we and others recently showed in cultured ECs (52, 53). Regardless of the underlying mechanism, by inducing kidney tissue hypoxia, capillary rarefaction can function as an upstream regulator of various defective responses (54). For example, Lovisa et al. showed an intriguing link between endothelial dysfunction and impaired tubular epithelial metabolism shifting from fatty acid oxidation to glycolysis (55).

Post-ischemic endothelial PHD inactivation augmented interstitial fibrosis and inflammation following kidney IRI, and these responses were associated with robust HIF activation. Based on our previous studies in which constitutive deletion of endothelial HIF-2 enhanced IRI-induced kidney fibrosis and inflammation (23), and pre-ischemic inactivation of endothelial PHD2 was protective (21), our new findings were unanticipated and raise new important questions. Does the timing of endothelial PHD inactivation in the setting of established kidney injury dictate the repair outcomes, does the non-physiologic endothelial HIF activation drive these unfavorable responses, or are HIF-independent effects of PHDs involved? These possibilities are of course not mutually exclusive. Pharmacological approaches of PHD inhibition applied prior to ischemic kidney injury have provided renoprotective effects, which though were not observed when HIF signaling was activated during the post-ischemic phase (56-60). Our study extends these findings on the timing of PHD inactivation being a critical factor in determining kidney injury responses by employing a genetic approach of inducible EC specific targeting. Furthermore, our system is not confounded by the potential off-target effects of PHD inhibitors on other 2-oxoglutarate-dependent dioxygenases. The observed suppression of fibrosis by inhibiting the HIF-1 regulated lactate transporter MCT4 supports the notion that glycolysis driven by persistent endothelial HIF-1 activation contributes to post-ischemic maladaptive kidney repair. Interestingly, Lim et al. reported that MCT4 downregulation alone significantly inhibits HIF transcriptional activity through a feed-back loop signaling (61), which could represent an additional mechanism underlying the renoprotective effects of syrosingopine. Nevertheless, it is plausible that non-HIF hydroxylation substrates of PHD could contribute to the observed outcomes by PHD inhibition. For example, PHD inhibition could reduce the hydroxylation of IKKβ derepressing NFkΒ signaling (62, 63), a response known to induce endothelial adhesion molecules binding and transmigration of leukocytes.

ECs are known to use glycolysis to meet > 80% of their ATP demands (64, 65), preserving oxygen for perivascular cells. Through scRNA-seq analysis, our study now unveiled that glycolysis can be further enhanced in post-ischemic renal ECs of *PHD^TiEC^* mice, which is in agreement with the metabolomic changes we recently reported in ECs treated with the pan-PHD inhibitor, dimethyloxalylglycine (DMOG) (53). Importantly, increased endothelial glycolysis was associated with pro-inflammatory changes that lead to enhanced macrophage infiltration. In accordance with our findings, recent studies reported induction of glycolysis in ECs subjected to the inflammatory mediators TNFA and lipopolysaccharide (42, 66), both known to induce HIF activation (67, 68). In this setting, among the significantly upregulated glycolytic genes, PFKFB3 was as identified a key regulator of this metabolic reprogramming. Notably, genetic or pharmacologic inactivation of PFKFB3 abrogated inflammation in various disease models such as LPS-induced inflammation and pulmonary hypertension demonstrating that PFKFB3-driven glycolysis drives endothelial pro-inflammatory pathways (66, 69). Consistent with the endothelial glycolytic signature observed in ECs in our scRNA-seq data, *Pfkfb3* showed a trend for upregulation, which though did not reach statistical significance probably due to limited power of our analysis. Importantly, by analyzing publicly available scRNA-seq data from kidneys of patients with severe AKI, we demonstrate that augmented endothelial glycolysis is observed in the human AKI endothelium along with hypoxia and inflammatory pathways such as IFN-γ, ΙFN-α and TNFA. Thus, our findings suggest a novel role for PHD/HIF in regulating immunometabolic phenotypes of kidney endothelium.

The HIF-1 induction of MCT4 allows the efficient export of lactic acid and enables glycolytic tumors to grow within an acidic microenvironment (70, 71). Furthermore, because MCT4-mediated lactic acid transport can regulate glycolytic flux, blocking MCT4 has been proposed as a strategy to suppress tumor growth in different cancers by disrupting the Warburg effect (72). Given that a) ECs resemble cancer cells in their preferential use of glycolysis b) MCT4 was significantly upregulated by endothelial PHD inactivation in the post-ischemic kidney, we postulated that MCT4 inhibition might promote adaptive kidney repair by suppressing endothelial glycolysis. To test this hypothesis, we treated *PHD^TiEC^* mice with syrosingopine starting on day 2 after uIRI and found that MCT4 inhibition remarkably reduced the tubular injury and development of fibrosis at day 14 post uIRI. To assess whether MCT4 inhibition had EC autonomous effects, HPAECs activated to a pro-inflammatory phenotype by hypoxia-reoxygenation and IL-1β, were treated with syrosyngopine or siMCT4. Both genetic silencing and pharmacologic MCT4 inhibition significantly suppressed the expression of EC-adhesion molecules reducing the adhesion of THP1 cells. Interestingly, inhibition of CD147, a broadly expressed transmembrane glycoporotein that is essential for MCT4 function (73), has been reported to reduce post-ischemic brain inflammatory cell infiltration by reduced NFkB activation in brain microvascular ECs (74). While this mechanism could explain our findings, a broader reprogramming of EC metabolism by MCT4 inhibition may be a key driver. For instance, Cluntun et al showed that MCT4 inhibition shifted glucose flux from lactate to citrate and TCA cycle in cardiomyocytes (75). Further studies are needed to investigate how metabolic reprogramming by MCT4 inhibition is linked to endothelial function.

In summary, our genetic studies demonstrate that post-ischemic endothelial PHD inactivation leads to HIF activation and maladaptive kidney repair, which can be reversed by MCT4 inhibition. Our findings merit further consideration of targeting the kidney endothelial hypoxia-driven glycolysis/MCT4 axis as a therapeutic strategy to halt the AKI to CKD transition.

## METHODS

### Generation of mice, genotyping, and animal procedures

The generation and genotyping of *Phd1* (*Egln2*), *Phd2* (*Egln1*), and *Phd3* (*Egln3*) floxed mice have been described previously (27, 28, 76). For inducible endothelial-specific inactivation of the floxed allele, we used the *Cdh5(PAC)-CreER^T2^*transgenic mouse line, provided by Dr. Ralph Adams (22). To examine recombination efficiency in the post-ischemic kidney, we generated *Cdh5(PAC)CreER^T2^;mT/mG* mice by crossing the ROSA26-ACTB-tdTomato,- EGFP reporter mice (JAX stock number 007576) to the *Cdh5(PAC)CreER^T2^* transgenic mouse line. Cre-mediated inactivation was induced by four intraperitoneal (IP) injections of tamoxifen dissolved in 10% ethanol in corn oil at 20 mg/mL (3 mg/mouse) given every other day. Mice were maintained in a specific pathogen-free facility on a 12-hour light/12-hour dark cycle. Littermates were randomly assigned to experimental groups.

Male mice 8-10 weeks of age underwent renal IRI surgery as previously described (77). Anesthesia was induced by IP injection of xylazine (10 mg/kg IP) and ketamine (90-120 mg/kg IP). After making a small midline abdominal incision, the left renal pedicle was occluded by applying a microaneurysm clamp for 25 minutes, while the right kidney was kept intact as an internal control in the unilateral IRI. For bilateral renal IRI, clamps were applied in both renal pedicles for 23 minutes. After the specified time, the clamps were removed, and reperfusion was visually confirmed. The incision site was closed with a 6-0 suture followed by the closure of the skin wound with Michel miniature clips. Throughout the surgical procedure, body temperature was monitored by a rectal probe and controlled with a heating pad at 37°C.

For in vivo MCT4 inhibition studies, mice were treated with syrosingopine (7.5 mg/kg) or vehicle starting on day 2 post IRI every other day until dissection on day 14 post IRI.

### Study approval

All animal studies were conducted in accordance with NIH guidelines for the use and care of live animal and were approved by the Institutional Animal Care and Use Committees at University of Kansas and at Northwestern University.

### DNA, RNA, and protein analyses

DNA and RNA were extracted and used for genomic or real-time PCR analysis, as previously described (14). Mouse and human primer sequences are listed in Supplemental Table 3. Real-time PCR was run in the QuantStudio 3 Real-Time PCR system (Applied Biosystems) using SYBR green or TaqMan PCR Master Mix. 18S rRNA was used for normalization.

Nuclear protein was extracted using NE-PER Nuclear and Cytoplasmic Extraction Reagents (Thermo Fisher Scientific), following the provided instructions. The extracted nuclear protein samples were then subjected to separation on an SDS-PAGE gel, transferred on a nitrocellulose membrane, and subsequently incubated with HIF-1α antibodies (Cayman) or HIF-2α antibodies (Novus Biologicals) at 4°C. Following an overnight incubation, the nitrocellulose membrane was washed and incubated with a secondary antibody (Novus Biologicals) for 90 minutes at 4°C. Chemiluminescent signal detection was performed using SuperSignal™ West Femto Chemiluminescent Substrate (Thermo Fisher Scientific), and membranes were imaged on the iBright Imaging System (Thermo Fisher Scientific).

### Histopathological and immunofluorescence analysis

For the histological examination, kidneys were fixed in 10% formalin buffer and paraffin blocks were prepared. To assess tubular damage and interstitial fibrosis, 5-µm transverse plane kidney sections were separately stained with hematoxylin and eosin (H&E) and Picro-Sirius red. Tubular injury was semi-quantitatively scored by determining the percentage of tubules in the corticomedullary junction that displayed necrosis, loss of brush border, cast formation, and tubular dilatation (0, unaffected; 1, 1%–25%; 2, 26%–50%; 3, 51%–75%; 4, 76%–100%) (78). Fibrosis was assessed by the percentage of Picro-Sirius red^+ve^ area determined with ImageJ software (http://rsbweb.nih.gov/ij). For tubular injury scoring and quantification of Picro-Sirius red^+ve^ area ζ10 and ζ5 random visual fields of corticomedullary region, respectively were analyzed per kidney section for each sample at a magnification of x200.

Immunofluorescence staining was performed using primary antibodies against PHD1 (Abcam; Catalog # ab113077), PHD2 (Novus Biologicals; Catalog # NB100-137), PHD3 (Novus Biologicals; Catalog # NB100-139), Endomucin (Abcam; Catalog # ab106100), F4/80 (Abcam; Catalog # ab6640), and MCT4 (Proteintech; Catalog # 22787-1-AP). Goat anti-rat Alexa Fluor® 488 (Invitrogen; Catalog# A11006), Goat anti-rabbit Alexa Fluor® 594 (Invitrogen; Catalog# A32740), and Goat anti-rabbit Alexa Fluor® 647 (Invitrogen, Catalog # A32733) were used as secondary antibodies. VECTASHIELD Vibrance Antifade Mounting Medium with DAPI (Vector Laboratories, Inc. Catalog # NC1601055) was used as mounting media. Images were captured by a fluorescence microscope (Nikon Ti2 widefield microscope) and analyzed using Fiji software (Image J). Percentage of Endomucin^+ve^area was determined with ImageJ software.

### BUN and GFR measurements

Serum BUN levels were measured using the QuantiChrom Urea Assay Kit (BioAssay Systems) following the manufacturer’s instructions. Transcutaneous GFR was measured via the transcutaneous clearance of FITC-Sinistrin using a miniaturized fluorescence detector (NIC-Kidney device) on day 14 post bIRI (79, 80).

### Preparation of single-cell suspension

Injured kidneys from *PHD^TiEC^* and *Cre^-^*mice were collected on day 14 post uIRI. Single-cell suspensions were prepared using Multi Tissue Dissociation Kit 2 (Miltenyi Biotec; Catalog # 130-110-203) following the instructions provided with the kit. Briefly, IR kidneys were cut into 6-8 pieces using single edge blades and placed in gentleMACS C tube with 5 ml dissociation buffer (4.8 mL of Buffer X, 50 µL of Enzyme P, 50 μL of Buffer Y, 100 µL of Enzyme D, and 20 µL of Enzyme A provided in Multi Tissue Dissociation Kit 2). GentleMACS C tubes were placed in a gentleMACS Dissociator (Miltenyi Biotec) and kidneys were dissociated using program Multi_E_01. Dissociated kidneys were incubated for 30 minutes at 37 °C with continuous rotation using the MACSmix Tube Rotator (Miltenyi Biotec). GentleMACS C tubes were then placed in the gentleMACS Dissociator and Multi_E_02 program was run. GentleMACS C tubes were detached and 10 mL neutralizing buffer (1X PBS with 2% fetal bovine serum) was added to stop dissociation reaction. Cell suspension was filtered through 100 µm and then 30 µm cell strainer. Cell suspension was centrifuged at 400 *g* for 10 minutes. Cell pellet was incubated with 1 mL of RBC lysis buffer on ice for 1 minute and after the addition of 10 mL neutralizing buffer, cells were recentrifuged at 400 *g* for 5 minutes. Cells were suspended in 10 mL neutralizing buffer and centrifuged again at 100 *g* for 5 minutes. Finally, cell pellet was suspended in 1X PBS with 0.04% BSA and filtered through 30 µm. Cell number and viability were analyzed using the Nexcelom Cellometer Auto 2000 with the AOPI fluorescent staining method. This method resulted in single-cell suspensions with > 80% viability.

### scRNA-seq library generation and sequencing

Sixteen thousand cells were loaded into the Chromium Controller (10X Genomics; Catalog # PN-120223) on a Chromium Next GEM Chip G (10X Genomics, Catalog # PN-1000120) and processed to generate single-cell gel beads in the emulsion (GEM) according to the manufacturer’s protocol. The cDNA and library were generated using the Chromium Next GEM Single Cell 3’ Reagent Kits v3.1 (10X Genomics; Catalog # PN-1000286) and single Index Kit T Set A (10X Genomics; Catalog # PN-1000213) according to the manufacturer’s manual. For the cell hashing library, it was constructed according to Biolegend’s published protocol on HTO library preparation guidelines with single index adapters. Quality control for the constructed library was performed by the Agilent Bioanalyzer High Sensitivity DNA Kit (Agilent Technologies; Catalog # 5067-4626), while Qubit DNA HS Assay Kit was used for qualitative and quantitative analysis. The multiplexed libraries were pooled and sequenced on an Illumina HiSeq 4000 sequencer with 2X50 paired-end kits using the following read lengths: 28 bp for Read1 for cell barcode and UMI, and 90 bp for Read2 for transcript. Raw sequencing data in base call format (.bcl) was demultiplexed using Cell Ranger from 10x Genomics, converting the raw data into FASTQ format. Cell Ranger was also used for alignment of the FASTQ files to the mouse reference genome (mm10) and to count the number of reads from each cell that align to each gene.

### scRNA-seq data analysis

The resulting matrix files, which summarize the alignment results, were imported into Seurat (Satija Lab, NYGC) for further analysis. In Seurat (v4.3.0), each individual sample was transformed to Seurat object. Only genes expressed in 3 or more cells, and cells expressing at least 200 genes, or more were used for further analysis. Further, cells with <500 unique molecular identifier (UMI) counts (cell fragments) and >100,000 UMI counts (potentially cell duplets) were excluded (81). Cells with mitochondrial gene percentage over 50% and low complexity cells such as red blood cells with <0.8 log10 genes per UMI counts were also excluded from the analysis (81). Merged data were normalized, scaled, and principal component analysis (PCA) was done. The top 20 principal components were chosen for neighbors and cell clustering performed with resolution of 0.8. The cell clusters were visualized in two-dimensional space by the Uniform Manifold Approximation and Projection (UMAP). Both genotypes showed similar clustering and cell populations in separate and overlapping DimPlot (82). To identify cell clusters, marker genes were assessed using “FindAllmarkers” function of Seurat with the setting of min.pct of 0.25 and logfc.threshold of 0.25. Based on marker analysis, one cluster predominately enriched with lncRNAs Gm26917 and Gm42418 associated with ribosomal RNA contamination was excluded (83, 84). The remaining 28 clusters were annotated based on the expression of top marker genes supported by published studies (Supplemental table 1). Differential gene expression analysis was performed, and genes with adjusted *P*<0.05, log_2_FC > 0.2 were considered significantly regulated. Gene set enrichment analysis was performed using GSEA software (v4.2.3) and by EnrichR (https://maayanlab.cloud/Enrichr/). Data were deposited in the NCBI’s Gene Expression Omnibus database (GEO GSE244064).

### Public scRNA-seq data analysis

#### Murine scRNA-seq data

To examine the expression of genes encoding the three HIF prolyl hydroxylase isoforms (*Egln1, Egln2 and Egln3)* in mouse kidney ECs, we extracted adult mouse kidney endothelial scRNA-seq data from the EC Atlas database created by the Carmeliet group (29). Data was processed and analyzed using standard Seurat’s workflow for data processing and normalization (https://satijalab.org/seurat/articles/pbmc3k_tutorial.html). Cell clusters were generated with 0.7 resolution and identified by analyzing the expression of EC marker genes (*Plvap*, *Igfbp3*, *S100a6*, *Sox17*, *Igfbp7*, *Aqp1*, *Pi16*, *Plat*, *Ehd3*, *Cyp4b*, and *Lpl*) (85).

#### Human scRNA-seq data

The expression of EGLNs genes in human kidneys ECs was assessed by extracting human kidney biopsy scRNA-seq datasets generated by Kidney Precision Medicine Project (KPMP) and available via Gene Expression Omnibus (GSE140989; Supplemental table 1) (30). Seurat files were created and combined following Seurat’s standard workflow. EC cell cluster was identified based on the expression of endothelial markers EMCN, CD34, PLVAP and EHD3. After extraction, EC were further clustered into 3 EC subclusters, which were identified based on the expression of different EC markers (*PLVAP*, *RGCC*, *MARCKS*, *SPARCL1*, *IGFBP7*, *TGFBR2*, *SOST*, *PLAT*, *EHD3*, *MEG3*, *SERPINE2*, *KCTD12*, *LTBP4*, *CLDN5*, and *IGF2*) (30).

#### Human snRNA-seq data from AKI and control kidney tissues

Human snRNA-seq datasets from AKI and control kidney tissues (8 AKI; 6 controls) were extracted from Gene Expression Omnibus (GSE210622) (39). Raw data was analyzed using Seurat’s best practice workflow for data integration using the reciprocal PCA approach (https://satijalab.org/seurat/articles/integration_rpca.html) with default parameters. Clusters were then analyzed for marker genes and EC cluster (*CD34*^+^ and *EMCN^+^*) was extracted, combined, and analyzed following Seurat’s best practice standard workflow for data integration and analysis (https://satijalab.org/seurat/articles/integration_introduction.html). Differential gene expression analysis was performed, and significantly upregulated genes (adjusted *P*<0.01, Log_2_FC>0.25) were subjected to hallmark analysis by EnrichR (https://maayanlab.cloud/Enrichr/). The expression levels of glycolytic genes in AKI and control kidney ECs were directly derived from the publicly accessible online interface (https://shiny.mdc-berlin.de/humAKI/) (39).

### Flow cytometry

Following perfusion with PBS, kidneys were harvested, minced, and incubated in dissociation solution (Multi Tissue Dissociation Kit 2; Miltenyi Biotec; Catalog # 130-110-203) for 30 minutes at 37°C. After dissociation, cells were passed through a 70- and 40- µm filter, and 10 mL of cold staining buffer (PBS + 2% fetal bovine serum) was added. Cells were centrifuged (400 *g* for 10 minutes), and cell pellets were resuspended in cold RBC lysis buffer, centrifuged, and resuspended in staining buffer. Following incubation with CD16/32 (Fc block), single-cell suspensions were labeled with different fluorophore-tagged antibodies. The following antibodies were used: CD16/32 (eBioscience™, Catalog # 14-0161-85), CD45 (clone 30-F1, Biolegend; Catalog # 103140), F4/80 (clone BM8, Biolegend; Catalog # 123110), CD11b (clone M1/70, Biolegend; Catalog # 101216), CD3 (clone 17A2, Biolegend; Catalog # 100210), Ly6C (clone HK1.4; Biolegend; Catalog # 128010), and Ly6G (BD biosciences; Catalog # 560600), and CD31 (clone 390, Invitrogen; Catalog # 17-0311-82).

### Cell culture

Human primary pulmonary artery ECs (HPAECs) were obtained from ATCC and grown on gelatin-coated dishes in Endothelial Cell Basal Medium-2 (Lonza, Catalog # CC-3156) supplemented with EGM-2 SingleQuots Supplements (Lonza, Catalog # CC-4176). MCT4 siRNA was purchased from Qiagen (FlexiTube GeneSolution GS9123 for SLC16A3; Catalog # 1027416). AllStars negative control siRNA (Qiagen) was used as control siRNA. HiPerFect transfection reagent (Qiagen, Catalog # 301707) was used for siRNA transfection experiments. For hypoxia experiments, MCT4siRNA transfected- or syrosingopine-treated HPAECs and their corresponding controls (AllStars negative control transfected and vehicle-treated, respectively) were incubated for 18 hours in a hypoxia chamber (Coy Laboratory Products) at 0.5% O_2,_ 37°C in a 5% CO_2_ humidified environment. During reoxygenation, HPAECs were treated with 1 ng/ml IL-1β (Millipore Sigma, Catalog# H6291) for 8 hours.

#### Endothelial-monocyte cell adhesion assay

For endothelial to monocyte adhesion assay, CellTracker™ Green CMFDA Dye (Invitrogen™, Catalog # C2925) labelled monocytes were added on HPAECs stimulated by hypoxia-reoxygenation and IL-1β. After 90 minutes, floating cells were removed, and fluorescent imaging was performed with quantification of the adhered monocytes using Fiji image J.

#### Lactate measurements

HPAECs were treated with vehicle (DMSO) or syrosingopine (5 μM) for 24 hours. Then, cell media were collected, and lactate levels were measured by a Lactate assay kit (Sigma, Catalog # MAK064) following the kit’s instruction and protocol.

### Statistical analysis

The data were expressed as mean values ± standard deviation of mean (SEM). Two-group comparison was performed by unpaired Student t-test with Welch’s correction. Multigroup comparison was performed by one-way analysis of variance (ANOVA) with Sidak’s multiple comparison test using GraphPad Prism 9 (GraphPad Software, La Jolla, CA). A value of *P* < 0.05 was considered statistically significant. Significance is shown by *P < 0.05, **P < 0.01, ***P < 0.001, ****P < 0.0001. ScRNA-seq and snRNA-seq analyses were performed using the program “R” (http://cran.r-project.org/), “RStudio” (https://posit.co/download/rstudio-desktop/) (https://posit.co/products/open-source/rstudio/) and Seurat (v4.3.0).

## AUTHOR CONTRIBUTIONS

RT and PPK conceived the study and designed the experiments; RT, RS, GR, GSB, SYA, JOS, MS, GC and PPK performed experiments; RT, RS, GR, GSB, SYA, JOS, MS, GC, VD, YZ, SEQ, ET, NSC and PPK analyzed and interpreted data; PPK and RT wrote the manuscript and all authors approved and commented on the manuscript.

## ACKNOWLEDGMENTS

This work was supported by National Institutes of Health (NIH) grants R01DK115850 and R01DK132672 (P.P.K.), and an American Heart Association (AHA) post-doctoral fellowship 23POST1020467 (R.T.). We acknowledge the Northwestern University George M. O’Brien Kidney Research Core Center (NU GoKidney), an NIH/NIDDK funded program (P30 DK114857) for their core services and support. Imaging work was performed at the Northwestern University Center for Advanced Microscopy generously supported by NCI CCSG P30 CA060553 awarded to the Robert H Lurie Comprehensive Cancer Center. Sc-RNA-seq was conducted at Northwestern University NUSeq Core Facility. Histology services were provided by the Northwestern University Research Histology and Phenotyping Laboratory, which is supported by NCI P30-CA060553 awarded to the Robert H Lurie Comprehensive Cancer Center. The funders had no role in study design, data collection and interpretation, or the decision to submit the work for publication. Graphical abstract was created with BioRender.com.

