## Supplemental Figures and Tables for "Post-ischemic inactivation of HIF prolyl hydroxylases in endothelium promotes maladaptive kidney repair by inducing glycolysis"

Tiwari et al.

The PDF file includes:
Supplemental Figures 1 to 9
Supplemental Tables 1 to 3

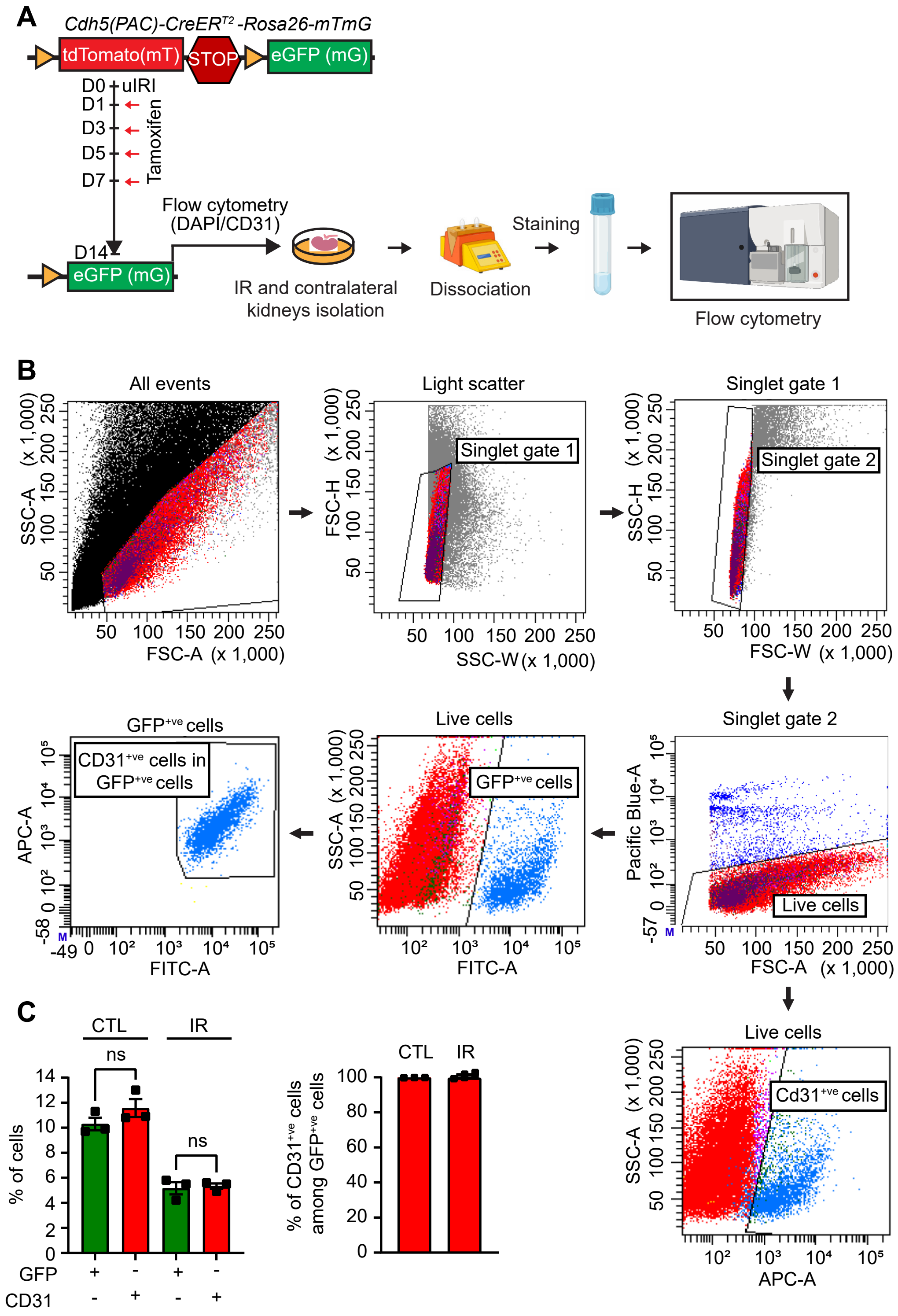


**Supplemental Figure 1, associated with Figure 1. Efficiency of *Cre*- recombination in *Cdh5(PAC)-CreER*$^{T2}$ transgenic mice following kidney IRI.** (**A**) By crossing *Cdh5(PAC)-CreER*$^{T2}$ mice with *ROSA26-ACTB-tdTomato,-EGFP (mTmG)* reporter mice, we generated *Cdh5(PAC)-CreER*$^{T2}$*; Rosa26-mTmG* mice, in which successful *Cre*- mediated excision is being indicated by GFP expression in endothelial cells (ECs). Mice were subjected to uIRI followed by tamoxifen administration starting at day 1 post uIRI to a total of 4 intraperitoneal (IP) injections given every other day. The degree of EC-specific recombination was assessed by FACs analysis in single-cell suspensions prepared by CTL and IR kidneys. (**B**) Flow cytometric gating strategy used to define recombined ECs. Staining for the endothelial marker CD31 was used to detect ECs, while GFP identified cells with tamoxifen induced expression of *Cdh5-CreER*$^{T2}$. (**C**) Shown are the percentages of GFP$^{+ve}$ and CD31$^{+ve}$ cells (left panel) as well as the percentage of CD31$^{+ve}$ cells among GFP$^{+ve}$ cells (right panel) in CTL and IR kidneys at day 14 post uIRI (n=3). For (**C**), statistics were determined by one-way ANOVA with Sidak correction for multiple comparisons. ns, not statistically significant. ECs, endothelial cells.

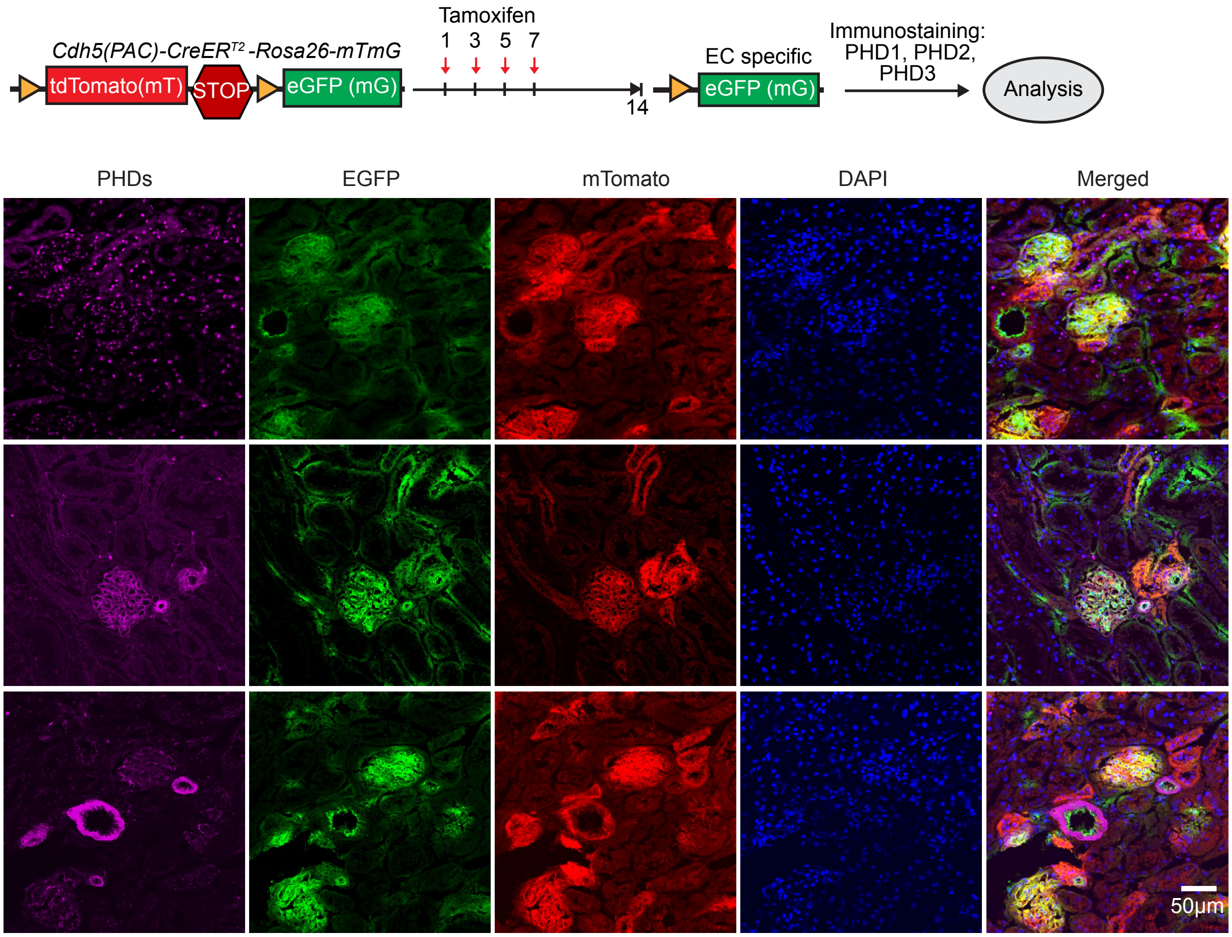


**Supplemental Figure 2, associated with Figure 2. Murine kidney endothelium shows compartment-specific differences in the expression of PHD1, PHD2 and PHD3.** Schematic view of experiment and representative images of immunofluorescence staining for PHD1, PHD2, PHD3 (magenta) and nuclear DAPI staining (blue) on kidney sections from *Cdh5(PAC)-CreER*$^{T2}$*-Rosa26mTmG* reporter mice after tamoxifen-induced recombination. Non recombined cells express membrane bound mTomato (red), whereas recombined cells express membrane-bound EGFP (green).

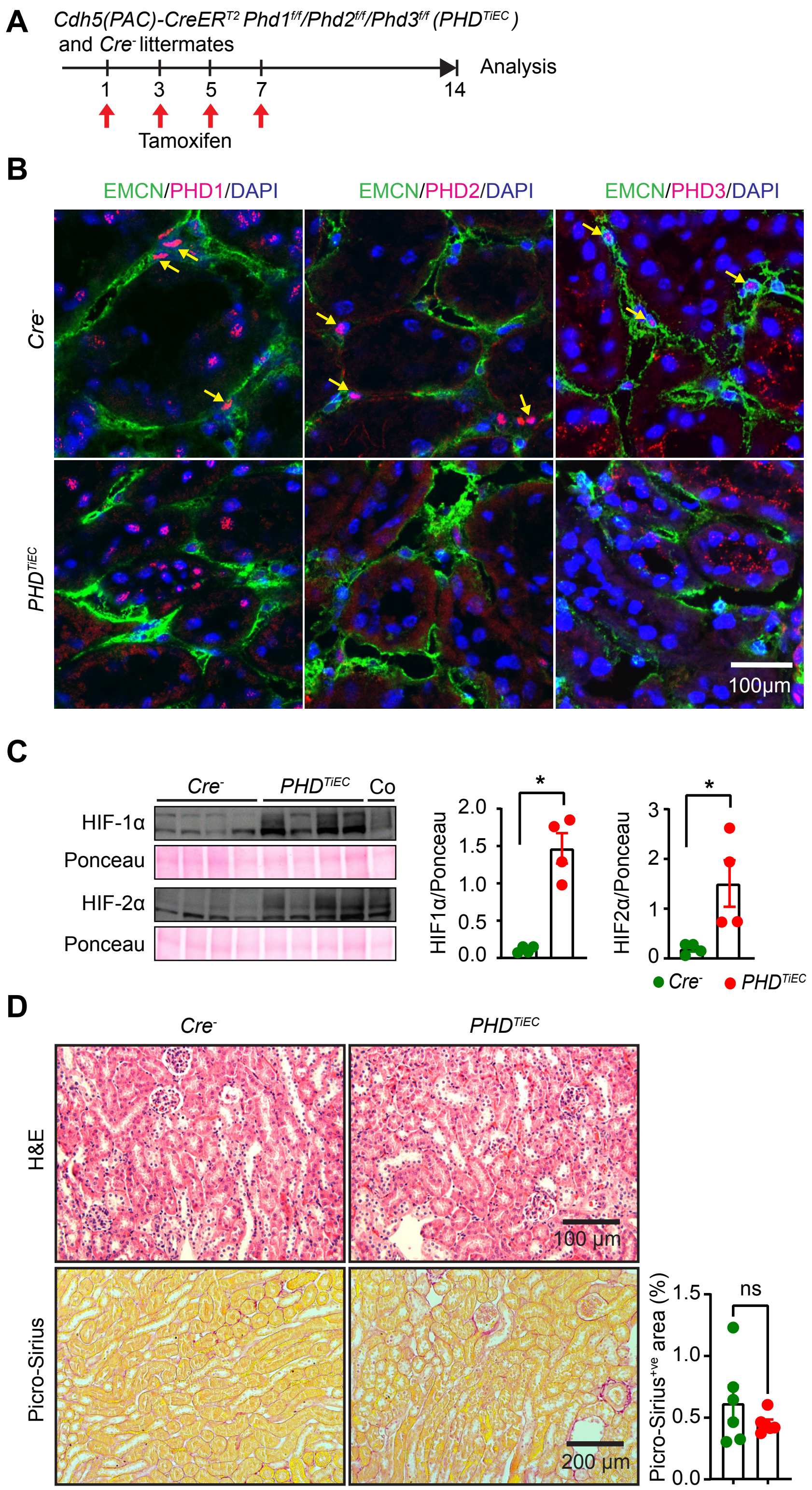


**Supplemental Figure 3, associated with Figure 3. Simultaneous acute inactivation of endothelial PHD1, 2, and 3 stabilizes HIF without significantly affecting kidney morphology.** (**A**) Schematic diagram depicting the tamoxifen administration regimen and the experimental strategy to obtain *PHD*$^{TiEC}$ mice and *Cre*$^{-}$ control mice. (**B**) Immunofluorescence staining for PHD1-3 (red), EMCN (green) and nuclear DAPI (blue) using kidney sections from *PHD*$^{TiEC}$ mice and *Cre*$^{-}$ controls collected 1 week after the last dose of tamoxifen. Scale bar, 100 µm. (**C**) Immunoblot analysis of HIF-1α and HIF-2α in kidney nuclear extracts isolated from *PHD*$^{TiEC}$ mice and *Cre*$^{-}$ littermates. Co, positive control. (**D**) Representative images of H&E and Picro-Sirius red stained kidney sections for the indicated genotypes. Right side graph shows semi-quantitative analysis of Picro-Sirius red$^{+ve}$ area for *PHD*$^{TiEC}$ mice and *Cre*$^{-}$ littermates 1 week after the last dose of tamoxifen (n=4-6). Scale bars indicate 100 µm and 200 µm for H&E and Picro-Sirius red images, respectively. Data are represented as mean ± SEM. Statistics were determined by unpaired t- test with Welch's correction. *, $P < 0.05$; ns, not statistically significant. EMCN, endomucin.

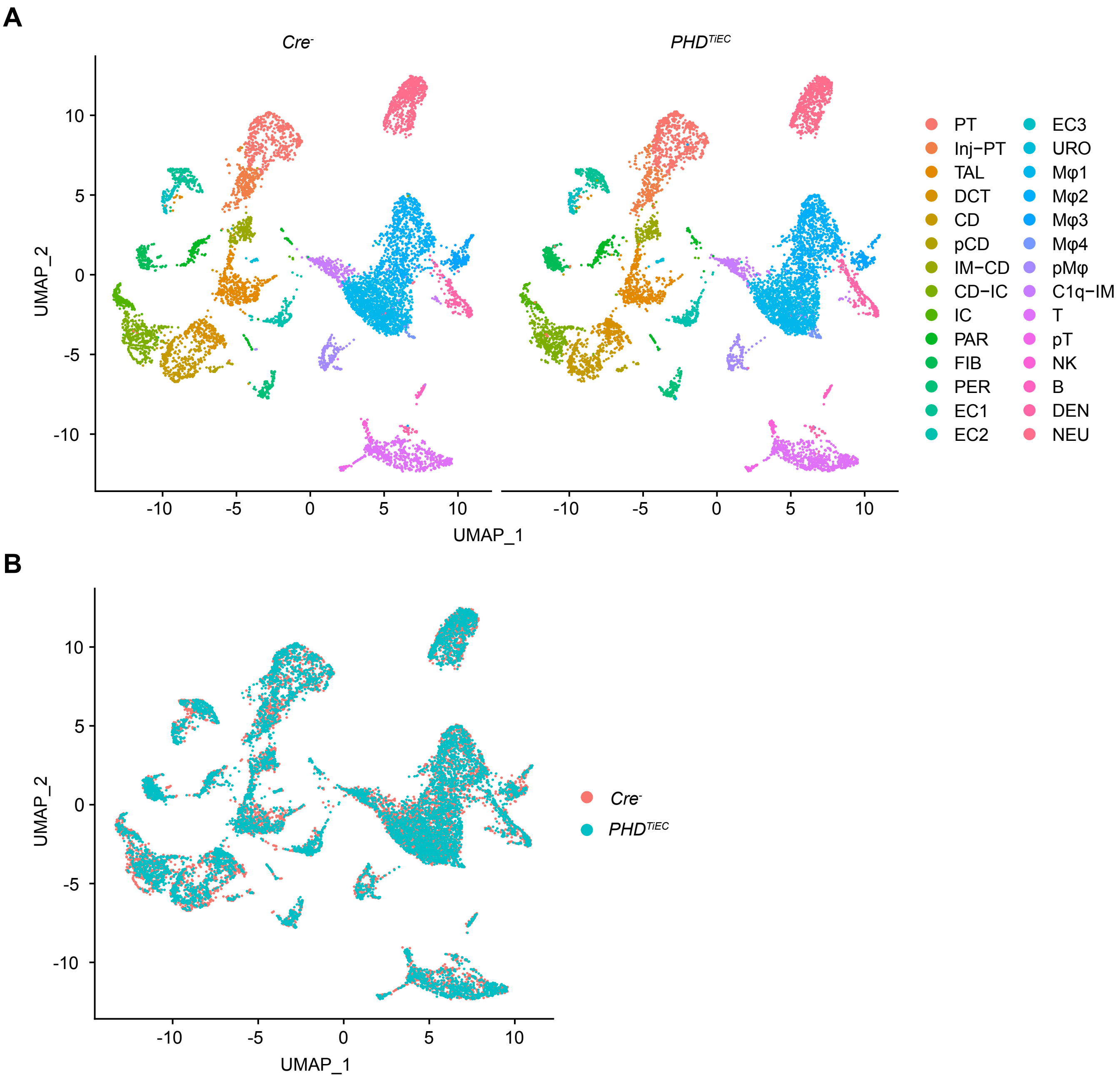


**Supplemental Figure 4, associated with Figure 4. scRNA-seq analysis showing similar cell populations in day 14 post-ischemic kidneys of *PHD$^{TiEC}$* and *Cre$^-$* mice.** (**A**) UMAP showing different cell clusters in day 14 post-ischemic kidneys from *PHD$^{TiEC}$* and *Cre$^-$* mice. (**B**) UMAP after overlaying samples of *Cre$^-$* and *PHD$^{TiEC}$* day 14 post-ischemic kidneys showing similar clustering.

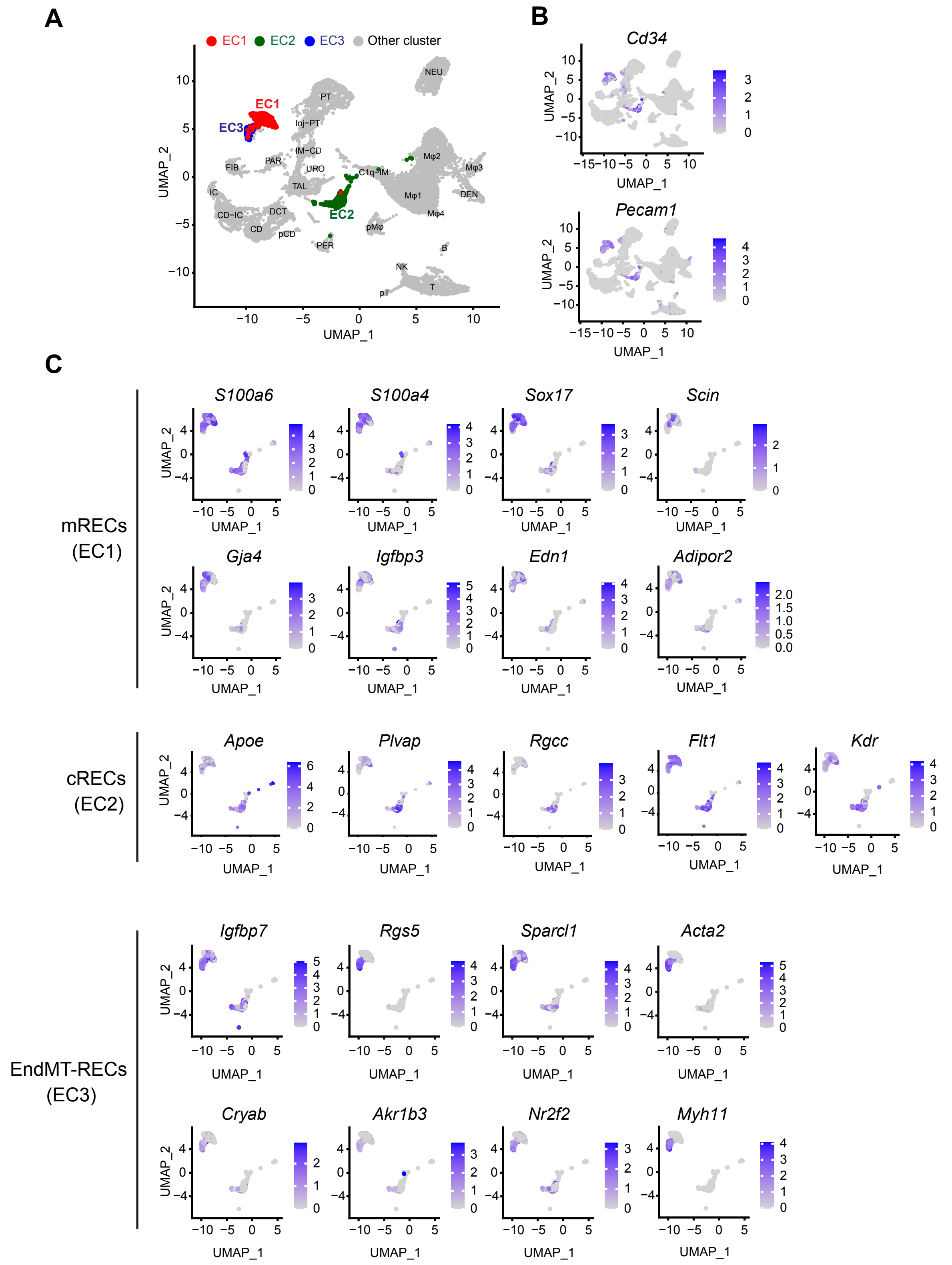


**Supplemental Figure 5, associated with Figure 5. Marker genes used to identify different EC clusters.** (**A**) UMAP plot with highlighted EC1 (red), EC2 (green) and EC3 (blue) cluster. (**B**) Feature plot showing expression of endothelial marker genes *Cd34* and *Pecam1* in EC1, EC2 and EC3 clusters. (**C**) Feature plot of EC marker genes used to identify mRECs (EC1), cRECs (EC2) and EndMT-RECs (EC3) clusters.

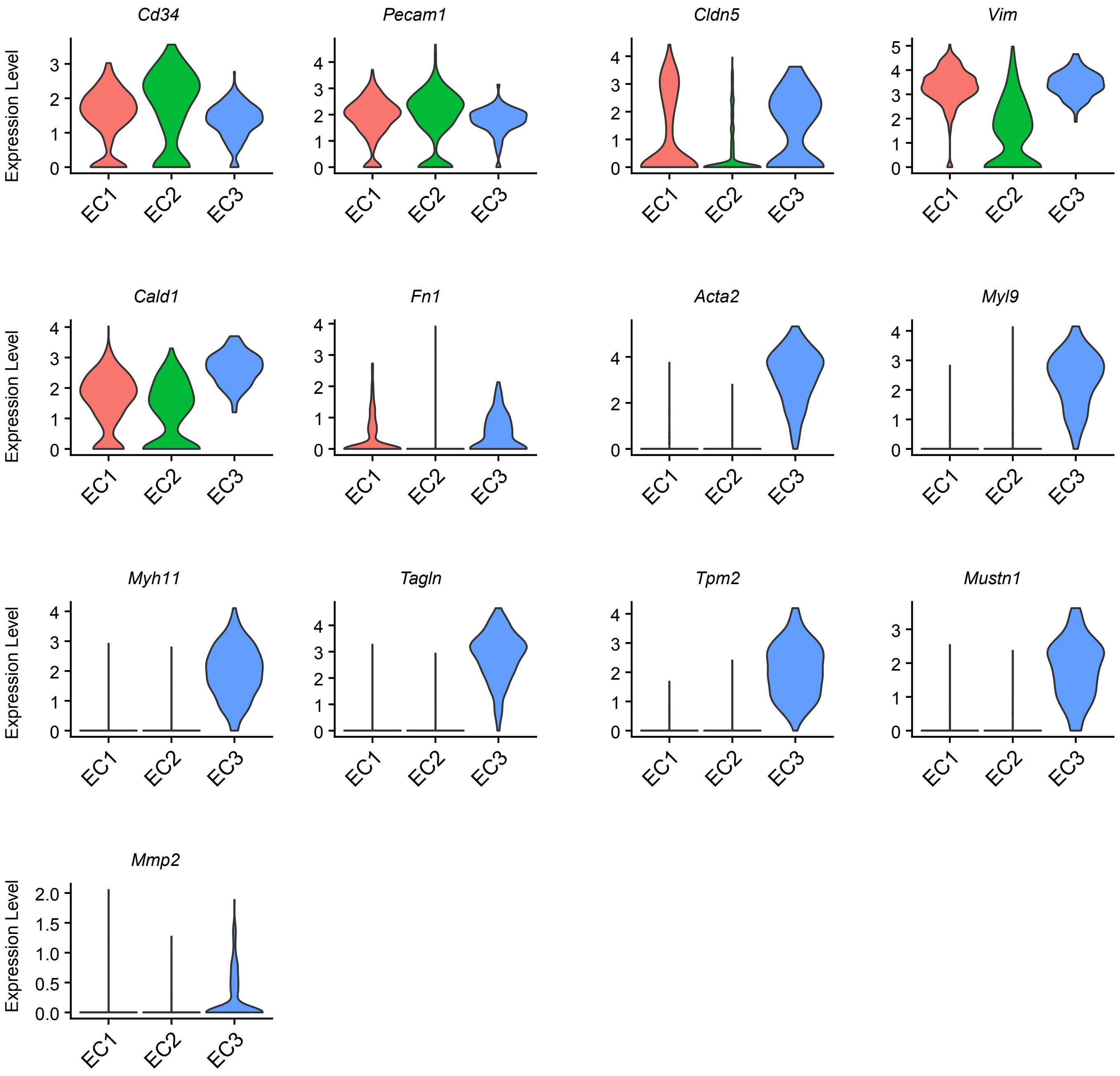


**Supplemental Figure 6, associated with Figure 5. Violin plots showing the expression of endothelial and mesenchymal markers in EC1, EC2 and EC3 clusters.** Violin plots show the expression of marker genes for ECs (*Cd34*, *Pecam1*, *Cldn5*) and mesenchymal cells (*Vim*, *Cald1*, *Fn1*, *Acta2*, *Myl9*, *Myh11*, *Tagln*, *Tpm2*, *Mustn1*, and *Mmp2*) in three EC clusters.

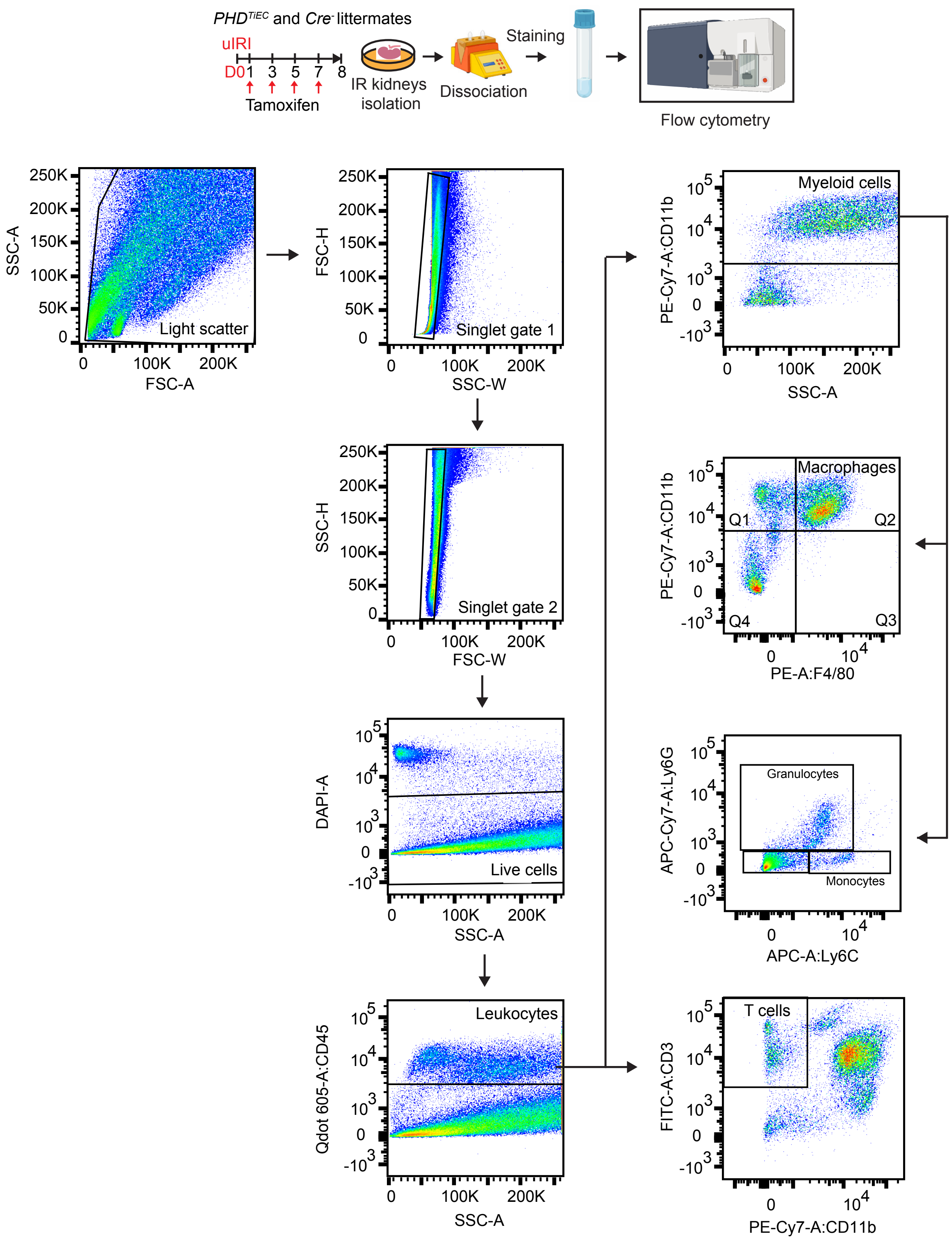


**Supplemental Figure 7, associated with Figure 6. Strategy for the analysis of immune cells by flow cytometry.** Schematic view of FACs experiment and gating strategy for the analysis of different immune cell populations in day 8 post-IRI kidneys from *PHD$^{TiEC}$* and *Cre*$^{-}$ mice. CD45$^{+}$ (leukocytes), CD45$^{+}$ CD3$^{+}$ (T cells), CD45$^{+}$ CD11b$^{+}$ (myeloid cells), CD45$^{+}$ CD11b$^{+}$ F4/80$^{+}$ (macrophages), CD45$^{+}$ CD11b$^{+}$ Ly6C$^{+}$ (monocytes), CD45$^{+}$ CD11b$^{+}$ Ly6G$^{+}$ (granulocytes) cells were measured.

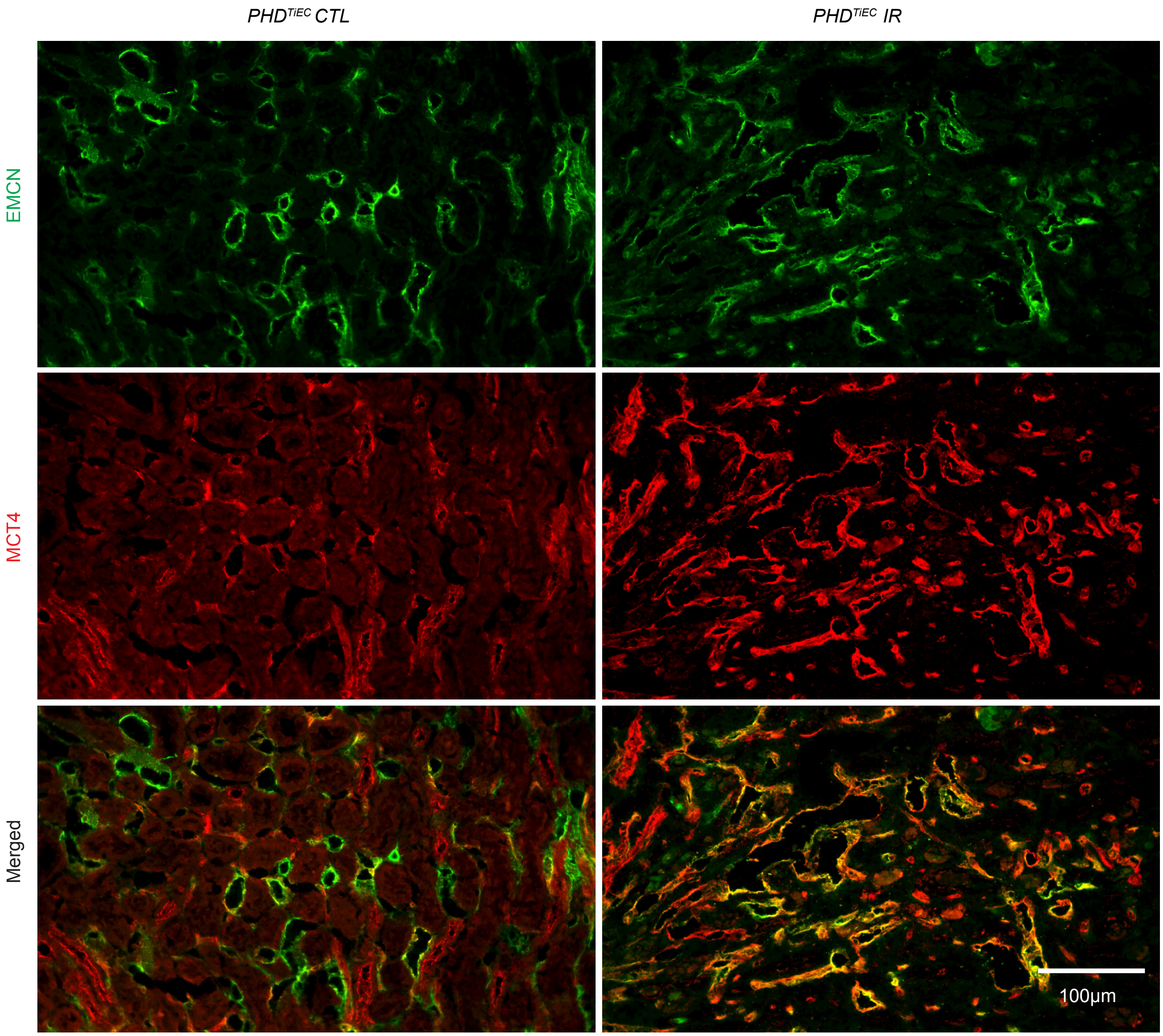


**Supplemental Figure 8, associated with Figure 7. MCT4 expression in contralateral and day 14 post-ischemic kidney of *PHD$^{TiEC}$* mouse .** Representative images of immunofluorescence staining for MCT4 (red) and EMCN (green) of CTL and IR kidneys on day 14 post-IRI from *PHD$^{TiEC}$* mice indicating increased expression of endothelial MCT4 following ischemic injury. Images were captured using Nikon a Ti2 Widefield fluorescence microscope. Scale bar,100 µm. CTL, contralateral kidney; IR, kidney subjected to uIRI.

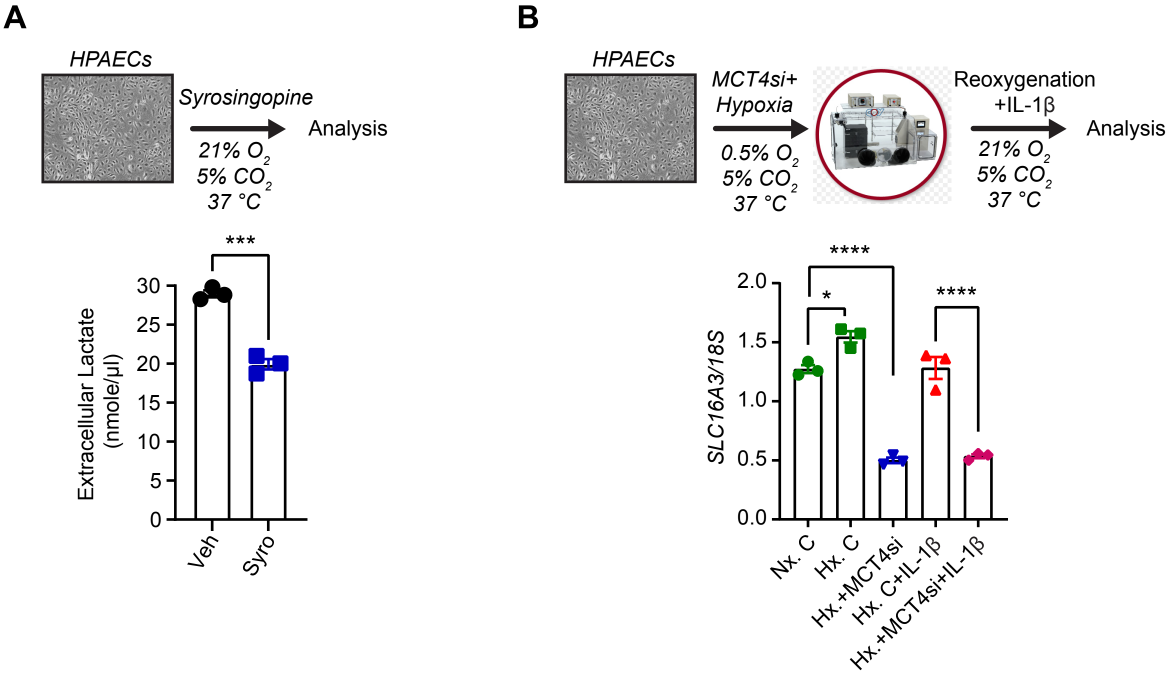


**Supplemental Figure 9, associated with Figure 8. Effect of syrosingopine on extracellular lactate levels and knockdown efficiency of MCT4 siRNA.** (**A**) HPAECs were treated for 24 hours with syrosingopine or vehicle (DMSO) and extracellular lactate levels were measured. (**B**) Experimental scheme for HPAECs subjected to 0.5% $O_2$ for 18 hours in the presence of MCT4 siRNA followed by reoxygenation for 8 hours in the presence of IL-1β (1 ng/ml). Graph showing *SLC16A3* (*MCT4*) mRNA levels in cells transfected with MCT4 siRNA compared to negative control siRNA transfected cells. All bars show mean ± SEM. For (**A**), unpaired t- test with Welch's correction was used. For (**B**), statistics were determined using one-way ANOVA with Sidak correction for multiple comparisons. *, $P$ <0.05; ***, $P$ <0.001; ****, $P$ < 0.0001. Veh, vehicle; Syro, syrosingopine; Nx, Normoxia; Hx, Hypoxia/Reoxygenation; C, negative control siRNA; MCT4si, MCT4 siRNA.

**Supplemental Table 1.** List of GEO numbers and details of samples which were used to assess the level of *Eglns* in human kidney ECs.

| Samples | GEO | Source name | Organism | Subject status | Tissue |
|---|---|---|---|---|---|
| 1 | GSM4191941 | Tumor-nephrectory | Homo sapiens | Normal/healthy | Adult kidney |
| 2 | GSM4191942 | Tumor-nephrectory | Homo sapiens | Normal/healthy | Adult kidney |
| 3 | GSM4191943 | Tumor-nephrectory | Homo sapiens | Normal/healthy | Adult kidney |
| 4 | GSM4191944 | Tumor-nephrectory | Homo sapiens | Normal/healthy | Adult kidney |
| 5 | GSM4191945 | Tumor-nephrectory | Homo sapiens | Normal/healthy | Adult kidney |
| 6 | GSM4191946 | Tumor-nephrectory | Homo sapiens | Normal/healthy | Adult kidney |
| 7 | GSM4191947 | Tumor-nephrectory | Homo sapiens | Normal/healthy | Adult kidney |
| 8 | GSM4191948 | Tumor-nephrectory | Homo sapiens | Normal/healthy | Adult kidney |
| 9 | GSM4191949 | Tumor-nephrectory | Homo sapiens | Normal/healthy | Adult kidney |
| 10 | GSM4191950 | Tumor-nephrectory | Homo sapiens | Normal/healthy | Adult kidney |
| 11 | GSM4191951 | Tumor-nephrectory | Homo sapiens | Normal/healthy | Adult kidney |
| 12 | GSM4191952 | Living donor | Homo sapiens | Normal/healthy | Adult kidney |
| 13 | GSM4191953 | Living donor | Homo sapiens | Normal/healthy | Adult kidney |

| | | | | | |
|---|---|---|---|---|---|
| 14 | GSM4191954 | Living donor | Homo sapiens | Normal/healthy | Adult kidney |
| 15 | GSM4191955 | Tumor-nephrectory | Homo sapiens | Normal/healthy | Adult kidney |
| 16 | GSM4191956 | Tumor-nephrectory | Homo sapiens | Normal/healthy | Adult kidney |
| 17 | GSM4191957 | Tumor-nephrectory | Homo sapiens | Normal/healthy | Adult kidney |
| 18 | GSM4191958 | Tumor-nephrectory | Homo sapiens | Normal/healthy | Adult kidney |
| 19 | GSM4191959 | Tumor-nephrectory | Homo sapiens | Normal/healthy | Adult kidney |
| 20 | GSM4191960 | Surveillance biopsy | Homo sapiens | Normal/healthy | Adult kidney |
| 21 | GSM4191961 | Surveillance biopsy | Homo sapiens | Normal/healthy | Adult kidney |
| 22 | GSM4191962 | Surveillance biopsy | Homo sapiens | Normal/healthy | Adult kidney |
| 23 | GSM4191963 | Surveillance biopsy | Homo sapiens | Normal/healthy | Adult kidney |
| 24 | GSM4191964 | Surveillance biopsy | Homo sapiens | Normal/healthy | Adult kidney |

**Supplemental Table 2.** List of used marker genes to identify different cell cluster of scRNA seq data of *Cre*[-] and *PHD*[TiEC] mice and their references.

| Marker genes | Cell cluster | Reference |
|---|---|---|
| *Pck1, Lrp2, Slc5a12, Slc34a1* | Proximal tubule (PT) | (1, 2) |
| *Havcr1* | Injured PT (Inj-PT) | (1) |
| *Umod, Slc12a1* | Thick ascending limb (TAL) | (1) |
| *Slc12a3* | Distal convoluted tubule (DCT) | (3) |
| *Fxyd4, Aqp2* | Collecting duct (CD) | (4) |
| *Slc14a2* | Inner medullary collecting duct (IM-CD) | (5) |
| *Atp6v1g3, Scl26a4* | Intercalated cells (IC) | (1, 6) |
| *Ncam1* | Parietal cells (PAR) | (7) |
| *Col1a2, Col1a2, Dcn* | Fibroblasts (FIB) | (2, 8, 9) |
| *Myh11, Acta2,* | Pericytes (PER)/smooth muscle cells | (9) |
| *Cd34, Pecam1, Igfbp3* | Endothelial cells (EC) | (1, 10) |
| *Upk1b* | Urothelial cells (URO) | (11) |
| *Mki67, Top2a* | Proliferating cells (p) | (1) (12) |
| *Ptprc* | Immune cells | (6) |
| *Itgam* | Myeloid cells | (12) |
| *Adgre1, Cx3cr1* | Macrophages (Mφ) | (1, 12) |
| *C1qa, C1qb, C1qc* | C1qa, C1qb and C1qc expressing immune cell (C1q-IM); potentially resident macrophage/monocyte | (13, 14) |
| *Trbc2, Skap1* | T cells | (14, 15) |
| *GZMA, GZMB* | Natural killer cells (NK cells) | (16) |
| *Ms4a1, Pax5, Bcl11a* | B cells | (15, 17-19) |
| *Flt3* | Dendritic cells (DC) | (20, 21) |
| *S100a8, S100a9* | Neutrophils (NEU) | (22) |

**Supplemental Table 3.** Primer sequences. Shown are sequences of primer sets used for the expression analysis of the indicated mouse and human genes by RT-PCR.

| Gene | Forward Primer | Reverse Primer |
|---|---|---|
| **Mouse** | | |
| *Loxl2* | 5′-GATCTTCAGCCCCGATGGA-3′ | 5′-CAAGGGTTGCTCTGGCTTGT-3′ |
| *Tgfb1* | 5′-TGGCGAGCCTTAGTTTGGA-3′ | 5′-TCGACATGGAGCTGGTGAAA-3′ |
| *Acta2* | 5′-CCTGACGCTGAAGTATCCGATAG-3′ | 5′-TTTTCCATGTCGTCCCAGTTG-3′ |
| *Kim1* | 5′-AAACCAGAGATTCCCACACG-3′ | 5′-GTCGTGGGTCTTCCTGTAGC-3′ |
| **Human** | | |
| *SLC16A3* | 5′-GGGTGGGAACCGTGTCATT-3′ | 5′-CTTGCGGCTTGGCTTCA-3′ |
| *VCAM1* | 5′-GCTTCAGGAGCTGAATACCC-3′ | 5′-AAGGATCACGACCATCTTCC-3′ |
| *ICAM1* | 5′-CCACAGTCACCTATGGCAAC-3′ | 5′-AGTGTCTCCTGGCTCTGGTT-3′ |
